# TestNet-PA: Sampling-Zero-Aware Inference of Microbial Presence-Absence Networks

**DOI:** 10.64898/2026.11.05.764020

**Authors:** Yicong Mao, Yi-Juan Hu

## Abstract

Microbial presence-absence networks, also referred to as co-occurrence networks, characterize patterns of coexistence and incompatibility among taxa. However, sequencing-based presence–absence analysis can be compromised by pervasive ‘sampling zeros’ in microbiome data, causing widely used methods such as Fisher’s exact test to identify spurious co-occurrence dependence. To our knowledge, no existing method is designed to account for sampling zeros in this setting. We introduce TestNet-PA, a method for testing pairwise microbial co-occurrence dependence. TestNet-PA models each observed zero as arising from a mixture of ‘structural’ and ‘sampling’ zeros, with the mixing probabilities governed by library size and relative abundance. Because the relative abundance underlying an observed zero is unobserved, we impute it by borrowing information from other samples. We then construct a quasi-likelihood and develop an EM algorithm to infer dependence between the underlying true presence–absence statuses. Extensive simulations based on real gut, vaginal, and oral microbiome data templates showed that TestNet-PA consistently controlled the FDR while maintaining good sensitivity. By contrast, Fisher’s exact test generally yielded inflated FDR. Analyses of the three real microbiome datasets further illustrated how accounting for sampling zeros can affect the inferred co-occurrence networks.

## Background

The human microbiome is increasingly recognized as an important determinant of health and disease, motivating growing interest in the inference of microbial networks. These networks characterize dependencies among taxa within a microbial community, including mutualism, competition, commensalism, and parasitism, and can help identify keystone taxa [1], delineate ecological guilds [2], or investigate disease-associated perturbations [3]. Throughout this article, we use the term ‘dependence’ broadly to encompass microbial relationships that may arise from either causal interactions or statistical associations.

To date, microbiome network studies [4–6] have largely focused on co-abundance networks, which characterize taxon–taxon dependencies in quantitative abundance measurements. This focus likely reflects, at least in part, the greater availability of statistical methods for co-abundance network inference [7–10], rather than greater scientific importance of abundance-based dependencies. For instance, we recently developed TestNet for inferring co-abundance networks based on centered-log-ratio transformed relative abundance data [10].

Co-occurrence networks capture patterns of coexistence and incompatibility among taxa at the presence–absence level, an important dimension of microbial community structure [11]. For example, one taxon may promote the presence of another by producing essential metabolites [12] or modulating host immune tone in ways that facilitate colonization [13]. Conversely, one taxon may inhibit another through competitive exclusion for essential resources [14] or by inducing host immune responses hostile to the other taxon [15]. We note that the term ‘co-occurrence’ has sometimes been used interchangeably with ‘co-abundance’ in the literature; here, we explicitly distinguish between the two.

More broadly, presence–absence analysis has received considerable attention as a robust and powerful alternative to relative-abundance analysis [16, 17]. By reducing quantitative count data to binary presence–absence statuses, it can sidestep several challenges inherent to microbiome data analysis. In particular, it is less affected by compositional constraints, avoids difficult choices involving data transformation and pseudocount imputation [18], and, more importantly, is more robust to experimental biases that distort relative abundance measurements [19]. Moreover, given the high sparsity of microbiome data, presence–absence analysis can itself be highly informative. In differential abundance analysis, presence–absence data performed comparably to relative-abundance data [20], while in disease-classification settings, presence–absence profiles alone achieved high predictive accuracy [21].

However, one critical challenge remains: the underlying presence–absence statuses are not always directly observed in sequencing-based experiments. Both 16S rRNA gene sequencing and metagenomic sequencing produce taxon count data with extremely high sparsity, often containing 60–95% zeros. Some zeros represent true taxon absence, commonly referred to as ‘structural zeros’, whereas many others arise because limited library size (sequencing depth) fails to detect taxa that are present at low abundance, giving rise to ‘sampling zeros’ [22]. Down-sampling studies suggest that sampling zeros may constitute a substantial fraction of observed zeros [23, 24]. If not properly accounted for, these sampling zeros can induce spurious co-occurrence dependencies, a manifestation of the ‘double-zero’ problem [25].

To illustrate this problem empirically, we collected three representative datasets from diverse body sites: a gut microbiome dataset from the Inflammatory Bowel Disease Multi-omics Database (IBDMDB) project [26], a vaginal microbiome dataset from the Multi-Omic Microbiome Study: Pregnancy Initiative (MOMS-PI) project [27], both from the Integrative Human Microbiome Project (HMP2) [28], and an oral microbiome dataset from the Oral Infections, Glucose Intolerance, and Insulin Resistance Study (ORIGINS) [29]. As shown in Figure 1(a), library sizes are limited and highly variable across samples within each study. The median library size was approximately 19,000 for IBDMDB, with an interquartile range (IQR) of [14,000, 23,000]; 47,000 for MOMS-PI, with an IQR of [25,000, 77,000]; and 133,000 for ORIGINS, with an IQR of [46,000, 250,000]. Figure 1(b) further shows a clear tendency for samples with larger library sizes to contain more observed taxa. This pattern is especially pronounced in the ORIGINS data.

**Figure 1:**
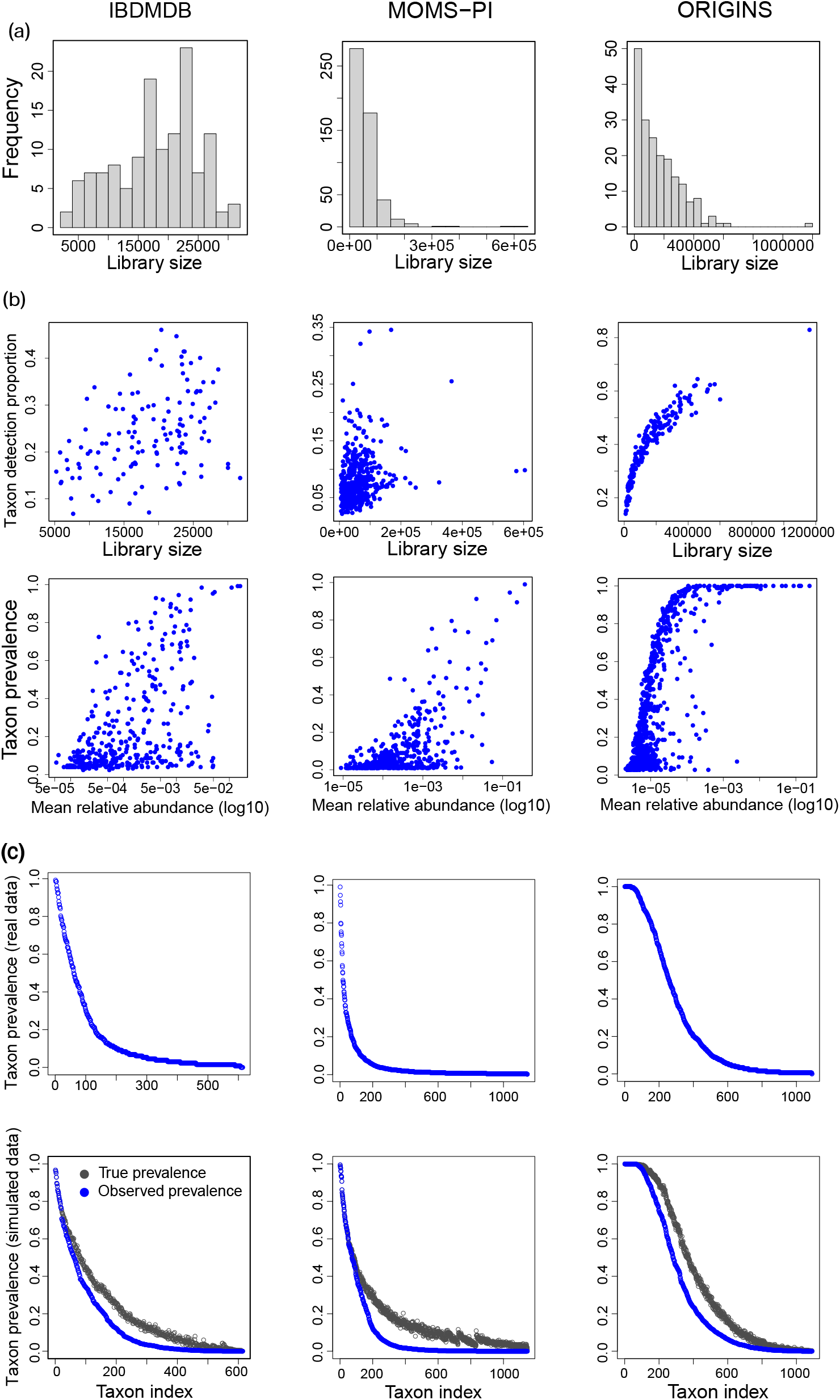
Characteristics of the three real microbiome datasets and corresponding simulated datasets. (a) Distributions of library sizes. (b) Taxon detection proportion per sample versus library size (upper panel), and taxon prevalence versus taxon mean relative abundance (lower panel), where the mean relative abundance was calculated using only nonzero relative abundances. (c) Taxon prevalence, sorted in descending order, for each real dataset (upper panel) and for a representative simulated replicate (lower panel).

Existing studies of microbial co-occurrence networks typically apply Fisher’s exact test or Pearson’s chi-squared test directly to observed zero–nonzero taxon count data, effectively treating all observed zeros as structural zeros [11, 30, 31]. Such analyses remain susceptible to library-size confounding and can produce many false-positive dependencies. Rarefaction, although commonly used to reduce library-size effects in differential abundance analysis [16, 17], offers little remedy in this setting. First, rarefaction substantially increases the frequency of sampling zeros and can result in essentially no power (results not shown). Second, forcing all samples to the same library size can induce spurious negative associations among sampling zeros through the compositional constraint, further distorting co-occurrence inference.

In this article, we introduce TestNet-PA for testing microbial co-occurrence dependencies while explicitly accounting for sampling zeros. We first formulate a latent-class model in which each observed zero arises from a mixture of structural and sampling zeros, with the mixing probabilities governed by library size and relative abundance and the dependence parameter of interest defined through the underlying true presence–absence statuses. We then address the challenge posed by unknown relative abundances and propose an imputation strategy. This formulation leads to a quasi-likelihood for inference, for which we establish theoretical validity. Through extensive simulation studies, we demonstrate both the advantages of TestNet-PA over Fisher’s exact test and its complementary value alongside state-of-the-art co-abundance methods. Finally, we apply TestNet-PA to the three aforementioned real datasets spanning the gut, vaginal, and oral microbiomes.

## Methods

### Latent-class model

For pairwise co-occurrence inference, we consider one pair of taxa, *j* and *k*, at a time. Let 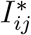 and 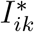 denote the *true, unobserved* presence–absence statuses of taxa *j* and *k*, respectively, in sample *i*, coded as 1 for presence and 0 for absence. Their joint status, 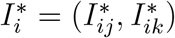, takes one of four values, ‘00’, ‘01’, ‘10’, and ‘11’, which define the four latent classes. Let *θ*_00_, *θ*_01_, *θ*_10_, and *θ*_11_ denote the corresponding class probabilities, which sum to 1. These probabilities encode the co-occurrence dependence of interest. We make this explicit by re-parameterizing the four *θ*’s in terms of (*π, α, β*). Specifically, 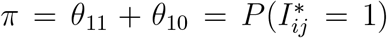 is the marginal prevalence of taxon *j*, while (*α, β*) are the intercept and slope in a logistic regression model for the conditional prevalence of taxon *k* given the status of taxon *j*:

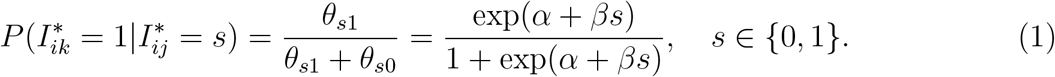

Under this parameterization, *β* = log[(*θ*_00_*θ*_11_)*/*(*θ*_01_*θ*_10_)] is the log odds ratio measuring co-occurrence dependence between taxa *j* and *k*. Thus, co-occurrence dependence is tested by *H*_0_ : *β* = 0, with (*π, α*) as nuisance parameters. For notational simplicity, the indices *j* and *k* are suppressed where no ambiguity arises.

We next link the true, unobserved presence–absence statuses to the observed sequencing data. Let *X*_*ij*_ denote the observed read count for taxon *j* in sample *i*. When 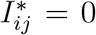, then necessarily *X*_*ij*_ = 0. When 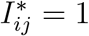, we assume that *X*_*ij*_ follows a count distribution with mean 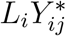, where *L*_*i*_ is the library size and 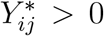 denotes the *potential* relative abundance of taxon *j* conditional on its presence 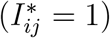. While *L*_*i*_ is known, 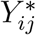 is generally unobserved. For now, we proceed as if 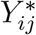 were also known and address its missingness later. Possible choices for the count distribution include the Poisson and negative binomial distributions. For our purpose, however, specifying the full count distribution is unnecessary; it suffices to model the sampling-zero probability *P* (*X*_*ij*_ = 0 | 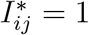).

Accordingly, rather than modeling *X*_*ij*_ directly, we work with the detection status, i.e., *observed* presence–absence status, *I*_*ij*_ = I(*X*_*ij*_ *>* 0), where I(·) denotes the indicator function. The sampling-zero probability, *P* (*I*_*ij*_ = 0 | 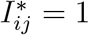), depends on the library size *L*_*i*_ and the relative abundance 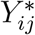 . As illustrated in Figure 1(b), taxa are more likely to be observed in samples with larger library sizes and at higher relative abundances. We therefore specify

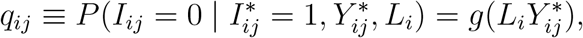

where *g*(.) is a decreasing, convex function mapping [0, ∞) to [0, 1]. This formulation accommodates a broad class of count-generating mechanisms. For example, under a Poisson model for *X*_*ij*_, *g*(*x*) = exp(−*x*); under a negative binomial model representing a Poisson–Gamma mixture, 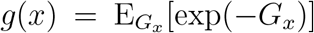, where *G*_*x*_ is a Gamma-distributed random variable with mean *x*. For our inference, we simply adopt *g*(*x*) = exp(−*x*) as a working model, without requiring it to be correctly specified. Thus, the proposed method is semiparametric in the sense that inference does not rely on a fully specified model for the observed data.

We can now formulate the latent-class model for the joint detection status of taxa *j* and *k*, denoted as *I*_*i*_ = (*I*_*ij*_, *I*_*ik*_). Modeling *I*_*i*_ as arising from a mixture over the latent classes, we obtain the following probabilities

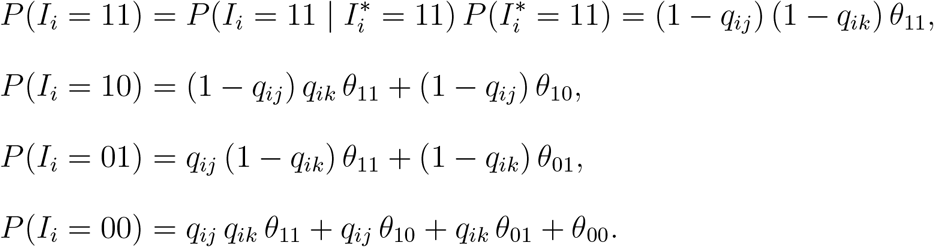

This formulation highlights the central role of the sampling-zero probabilities in distinguishing structural zeros from sampling zeros. Because each *q*_*ij*_ depends on library size, an observed zero arising in a deeply sequenced sample is more likely to reflect a true absence, whereas one arising in a shallowly sequenced sample is more likely to be a sampling zero. Although *q*_*ij*_ also depends on the taxon-specific relative abundance 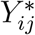, library size is the source of cross-taxon confounding because the same *L*_*i*_ enters the sampling-zero mechanism for both taxa.

### Likelihood with known relative abundances

Suppose that the relative abundances 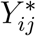 are known, so that the sampling-zero probabilities *q*_*ij*_ are also known. Then, the log-likelihood for Θ = (*θ*_00_, *θ*_01_, *θ*_10_), or equivalently Θ = (*π, α, β*), based on the detection statuses *I*_*i*_ from *n* samples is 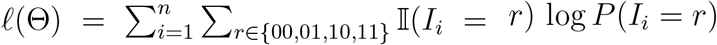. After dropping terms that depend only on the known *q*_*ij*_’s, this becomes

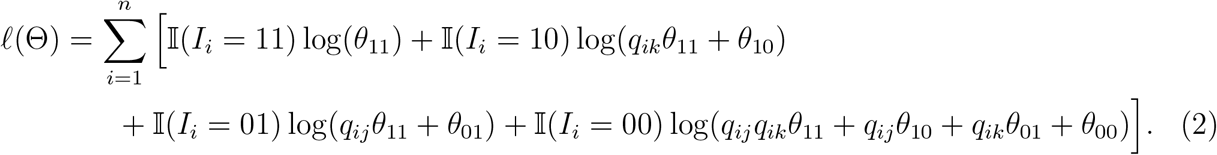

The difficulty is that *q*_*ij*_ or *q*_*ik*_ is needed precisely when the corresponding observed status is zero, *I*_*ij*_ = 0 or *I*_*ik*_ = 0, which is also when the associated relative abundance, 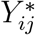 or 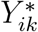, is particularly difficult to estimate, as discussed in the next section. Consequently, the likelihood in (2) is not directly available from the observed data, motivating an approach to recover or approximate the missing 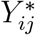’s.

### Relative abundance imputation

When a taxon is truly present 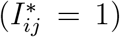 and detected (*I*_*ij*_ = 1, equivalently *X*_*ij*_ *>* 0), its relative abundance 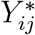 is estimable from the empirical relative abundance, *X*_*ij*_*/L*_*i*_. In contrast, when a taxon is undetected (*I*_*ij*_ = 0, equivalently *X*_*ij*_ = 0), the observed zero may be a sampling zero, in which case 1*/L*_*i*_ provides at best an upper-bound estimate for 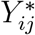, or it may be a structural zero, in which case the observation provides no information about 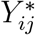 . Because these two cases cannot be distinguished from the observed data, direct estimation of 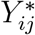 is particularly challenging when *I*_*ij*_ = 0.

We adopt an imputation strategy that borrows information about 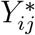 from other samples. We assume that, had the taxon been present in a sample where it is undetected, its potential relative abundance would have the same mean as in the population of samples. This motivates imputing 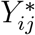 for *I*_*ij*_ = 0 by 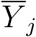, the average empirical relative abundance among samples in which taxon *j* is detected. Because detection preferentially occurs at higher relative abundances, 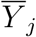 tends to overestimate the mean potential relative abundance; low-abundance realizations are disproportionately excluded from the detected samples. As shown in our numerical evaluations, however, this overestimation has little impact on inference.

### Quasi-likelihood

Plugging 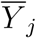 into the sampling-zero probability *q*_*ij*_ for observations with *I*_*ij*_ = 0 gives the working sampling-zero probability, 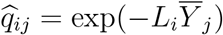, which is then substituted for *q*_*ij*_ in *l*(Θ), yielding the following quasi-likelihood:

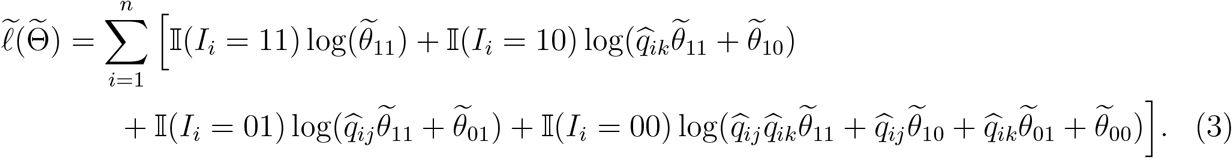

We refer to 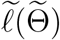as a quasi-likelihood because it is no longer a genuine likelihood but rather a misspecified working likelihood. Because 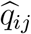enters the quasi-likelihood only when *I*_*ij*_ = 0, nonzero read counts contribute to the quasi-likelihood indirectly, through 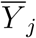 Compared with Fisher’s exact test, which relies solely on the binary detection statuses *I*_*ij*_, our quasi-likelihood additionally exploits information in the library sizes and nonzero read counts through 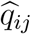.

Note that we introduce a distinct set of parameters targeted by the quasi-likelihood, 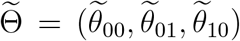and 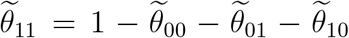, to distinguish them from the parameters Θ = (*θ*_00_, *θ*_01_, *θ*_10_) and *θ*_11_ in the true likelihood *l*(Θ), as 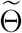 and Θ need not coincide under mis-specification. Analogous to Θ, we reparameterize 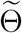 in terms of 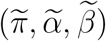, where 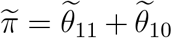,

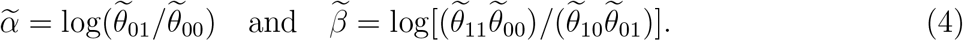

Importantly, despite the misspecification, we show next that the null is preserved, with 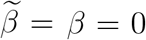 under no co-occurrence dependence, while under the alternative, 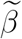 retains the sign of *β* but is attenuated toward zero.

### Quasi-likelihood as the true likelihood with omitted random effects

To establish the relationship between the quasi-likelihood 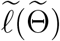 and the true likelihood *l*(Θ), we rewrite *l*(Θ) in terms of the working sampling-zero probabilities used in 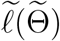. Up to an additive constant *C*, this yields

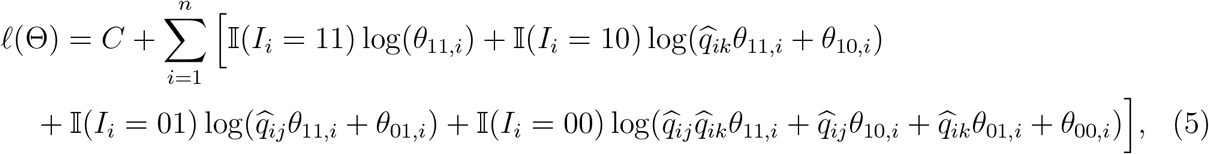

where the new class weights are now sample specific:

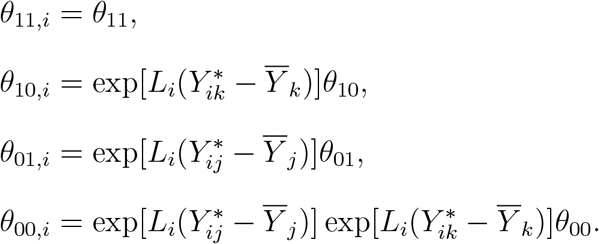

Note that these weights are not normalized to sum to one.

Comparing the new class weights with the original class probabilities Θ reveals three useful properties. First, it follows immediately that

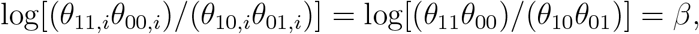

so the log odds ratio characterizing taxon–taxon dependence is common across samples and remains equal to the true value *β*. Second, log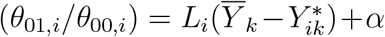, since log(*θ*_01_*/θ*_00_) = *α*, so 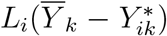 acts as a sample-specific random effect. Third, log 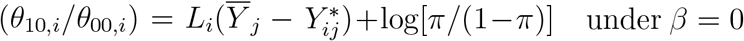, in which case log(*θ*_10_*/θ*_00_) = log[*π/*(1−*π*)], so 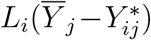 has the same interpretation of a random effect. Taken together, comparing the quasi-likelihood (3) with the true likelihood (5), the quasi-likelihood can be viewed as omitting sample-specific random effects from the true likelihood, while preserving the log odds ratio.

Ignoring sample-specific random intercepts is known to attenuate regression coefficients in logistic models [32]. Attenuation of covariate effects has also been extensively studied in the errors-in-variables literature [33, 34], which is closely related to random-effects formulations. We establish the corresponding result for our latent-class model in the Supplementary Ma-terials: under the assumption that the random effects have mean zero, the quasi-likelihood parameter 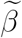 is attenuated toward zero relative to 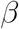 when *β* ≠ 0, while the null is preserved, with 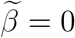 when *β* = 0.

### Quasi-likelihood-based inference

We estimate 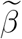 by maximizing the quasi-likelihood 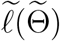 in (3) through an expectation-maximization (EM) algorithm (see the Supplementary Materials for full details). When one or both taxa in a pair have low prevalence, the presence status of one taxon may (nearly) perfectly predict that of the other, giving rise to complete or near separation. Such separation can lead to severe bias or even infinite parameter estimates. We therefore incorporate a Firth-like bias-reducing correction [35] (Supplementary Materials) into the M-step of the EM algorithm. Let 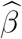 denote the resulting estimate of 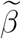.

In general, under likelihood misspecification, the variance estimator of 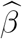 takes a sandwich form. In our setting, however, we found the sandwich estimator to be unstable in small to moderate samples (Figure S1). We therefore use the inverse observed information from the quasi-likelihood, computed using the Louis formula [36], as a more stable variance estimator. This choice is motivated by the fact that, under the null hypothesis (*β* = 0), the Louis estimator agrees with the sandwich estimator to first order in the random effects (Supplementary Materials). Its accuracy is further supported by our simulation results.

For each taxon pair, we then construct a Wald statistic and obtain the corresponding *p*-value from its asymptotic standard normal distribution. The *p*-values are subsequently adjusted using the Benjamini–Hochberg (BH) procedure [37] to control the false discovery rate and identify significant taxon pairs.

## Results

### Main simulation settings

Our main simulation studies were conducted using the three microbiome datasets described above as templates, which span a broad range of microbiome data characteristics. Specifically, following the preprocessing procedures detailed in the Supplementary Materials, the IBDMDB gut microbiome dataset comprised 127 samples and 614 genera, the MOMS-PI vaginal microbiome dataset comprised 502 samples and 1,146 species, and the ORIGINS oral microbiome dataset comprised 183 samples and 1,091 genera (Table 1). In addition to the substantial differences in their library size distributions noted in the Introduction, these datasets also differ markedly in sparsity. The vaginal dataset is the sparsest, with 95% zeros, followed by the gut dataset with 85% zeros and the oral dataset with 62% zeros.

**Table 1:** Dependent taxon pairs identified by co-occurrence or co-abundance analysis in the real datasets.

| Project | Body site | *n* | *L*<sup>†</sup> | *J* |  | Number of pairs | Method | Detected pairs | Positive pairs |
|---|---|---|---|---|---|---|---|---|---|
|  |  |  |  | Initial | Post-filter |  |  |  |  |
| IBDMDB (16S) | Gut | 127 | 20000 | 614 | 277 | 38226 | TestNet-PA | 870 | 860 |
|  |  |  |  |  |  |  | Fisher | 1716 | 1693 |
|  |  |  |  |  |  |  | TestNet | 1811 | 1684 |
|  |  |  |  |  |  |  | Spearman-RA | 2937 | 2865 |
|  |  |  |  |  |  |  | SparCC | 291 | 243 |
|  |  |  |  |  |  |  | SECOM-linear | 3007 | 1457 |
|  |  |  |  |  |  |  | SECOM-nonlinear | 0 | 0 |
| MOMS-PI (16S) | Vaginal | 502 | 47000 | 1146 | 201 | 20100 | TestNet-PA | 2741 | 2507 |
|  |  |  |  |  |  |  | Fisher | 2751 | 2464 |
|  |  |  |  |  |  |  | TestNet | 3424 | 2726 |
|  |  |  |  |  |  |  | Spearman-RA | 3635 | 3189 |
|  |  |  |  |  |  |  | SparCC | 206 | 181 |
|  |  |  |  |  |  |  | SECOM-linear | 1032 | 604 |
|  |  |  |  |  |  |  | SECOM-nonlinear | 0 | 0 |
| ORIGINS (MGS) | Oral | 183 | 133000 | 1091 | 384 | 73536 | TestNet-PA | 211 | 211 |
|  |  |  |  |  |  |  | Fisher | 14801 | 14801 |
|  |  |  |  |  |  |  | TestNet | 8484 | 8400 |
|  |  |  |  |  |  |  | Spearman-RA | 10241 | 10186 |
|  |  |  |  |  |  |  | SparCC | 200 | 200 |
|  |  |  |  |  |  |  | SECOM-linear | 2330 | 1613 |
|  |  |  |  |  |  |  | SECOM-nonlinear | 0 | 0 |

Note: Co-occurrence methods are listed above the short horizontal line, and co-abundance methods below it. <sup>†</sup>: approximate median library size; PA: presence–absence; RA: relative abundance; MGS: metagenomic sequencing; *n*: sample size; *J*: number of taxa. The filter excluded taxa with prevalence < 5% or > 80%. Results are reported at the 1% nominal FDR level, except for SparCC, for which an absolute estimated correlation of 0.3 was used as the significance threshold.

To generate presence–absence statuses with co-occurrence dependence, we used a multivariate normal (MVN) model followed by truncation. Specifically, for each sample *i*, we first generated a set of latent continuous variables as

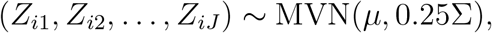

where *J* was set to the number of taxa in the corresponding data template, *µ* was a mean vector specified below, and Σ = {*σ*_*jk*_} was a prespecified covariance matrix. The factor 0.25, equivalent to scaling the standard deviations by 0.5, was used to regulate the overall variability of the latent variables. Following [10], we considered the following six covariance structures:

- (Identity) Σ = *I*_*J*_, corresponding to the global null setting.
- (AR1+) *σ*_*jk*_ = 0.5 if |*j* − *k*| = 1 and 0 otherwise, with *σ*_*jj*_ = 1.
- (AR1) *σ*_*jk*_ = ±0.5 if |*j* −*k*| = 1 and 0 otherwise, where the sign is assigned independently with equal probability, with *σ*_*jj*_ = 1.
- (AR4) *σ*_*jk*_ = ±0.1 if |*j* − *k*| = 4, ±0.5 if |*j* − *k*| = 2 or 3, ±1 if |*j* − *k*| = 1, and 0 otherwise. The signs are assigned independently with equal probability. The diagonal elements *σ*_*jj*_ are chosen to be sufficiently large to ensure that Σ is positive definite.
- (Block) The *J* taxa are divided as evenly as possible into 10 sets. Within each set of size *s*, the covariance matrix is the unit-diagonal normalization of ΛΛ^⊤^ + *I*_*s*_, where Λ ∈ ℝ ^*s*×1^ has entries with magnitudes sampled from Unif[0.7, 0.95] and signs assigned independently with equal probability. Covariances between taxa belonging to different sets are set to zero.
- (Sparse) Σ = diag(*D*_1_, *D*_2_), where *D*_1_ = *I*_*J*−*s*_ with *s* = 350 if *J* = 1091 and *s* = 300 otherwise, and *D*_2_ is the unit-diagonal normalization of ΛΛ^⊤^ + 0.1*I*_*s*_. Here, Λ ∈ ℝ ^*s*×3^ is a sparse matrix with 20% nonzero entries. The magnitudes of the nonzero entries are sampled from Unif[0.5, 0.9], with signs assigned independently with equal probability.

We sampled the elements of the mean vector *µ* = {*µ*_*j*_} from template-specific distributions chosen to reproduce the taxon prevalence patterns observed in the corresponding data templates. Specifically, we sampled *µ*_*j*_ ∼ 2.8 + 10 × Beta(10, 50) for the gut-based simulations, *µ*_*j*_ ∼ 4 + 15 ×Beta(1.5, 50) for the vaginal-based simulations, and *µ*_*j*_ ∼ 3.5 + 10 ×Beta(1.5, 10) for the oral-based simulations. The true presence–absence statuses were then obtained by truncating the latent *Z*_*ij*_’s at 5:

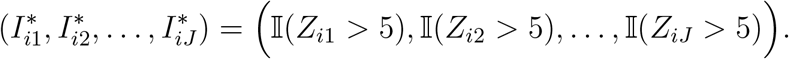

Under this construction, Σ encodes the co-occurrence dependence structure: *σ*_*jk*_ = 0 indicates no co-occurrence dependence, whereas *σ*_*jk*_ *>* 0 and *σ*_*jk*_ *<* 0 indicate positive and negative dependence, corresponding to coexistence and incompatibility, respectively.

Independently of the true presence–absence statuses 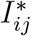, we generated the *potential* relative abundances 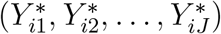 from a Dirichlet(*v, ϕ*) distribution. The mean vector *v* was set to the mean relative abundances estimated from the corresponding data template, and the overdispersion parameter was set to *ϕ* = 10^−5^ (a smaller value means less variation) to prevent rare taxa from exhibiting excessive variability and becoming spuriously abundant in some samples. Finally, for samples in which 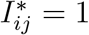, we generated the read count *X*_*ij*_ for taxon *j* from a negative-binomial distribution with mean 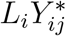 and overdispersion parameter 5 (with a larger value indicating less variation), allowing for moderate count-level overdispersion conditional on the potential relative abundances. The library size *L*_*i*_ was sampled from the empirical distribution of library sizes in the corresponding data template. When 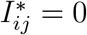, we set *X*_*ij*_ = 0. We repeated this procedure to generate *n* = 100 to 1, 000 samples in each replicate.

Figure 1(c) confirms that the simulated datasets closely reproduce the taxon-prevalence patterns observed in the corresponding data templates. The discrepancies between the observed and true prevalence reflect the occurrence of sampling zeros. The proportions of sampling zeros generated in our simulations may be conservative relative to those encountered in practice [23, 24], potentially giving Fisher’s exact test a relative advantage over TestNet-PA.

### Simulation settings for comparison with co-abundance methods

Although our main simulation settings closely reproduce the characteristics of the three data templates, they are not well suited for comparisons with co-abundance methods, which target quantitative dependence but may also detect qualitative dependence. In particular, the Dirichlet sampling step may induce co-abundance dependence among null taxon pairs with no co-occurrence dependence, potentially inflating the FDR of co-abundance methods. To enable a fairer comparison, we considered a second set of simulation settings adapted from the TestNet simulation framework [10]. In this framework, absolute abundances are first generated independently across taxa to avoid spurious co-abundance dependence, allowing us to introduce and evaluate co-occurrence dependence in isolation.

Specifically, we first generated the underlying log-scale absolute abundances (*W*_*i*1_, *W*_*i*2_, …, *W*_*iJ*_ ) from MVN(*w, I*_*J*_ ), where the elements of *w* were independently sampled from Unif[0, 10].

The identity covariance matrix *I*_*J*_ ensures independence of the absolute abundances across taxa. The *potential* relative abundances were then obtained by normalizing 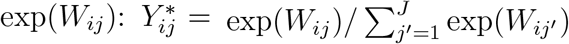. We next generated the true presence–absence statuses 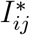, inde-pendently of the absolute abundances, using the same procedure as in the main simulation settings, but with *µ* = 3.8 + 10 × Beta(5, 50) to yield approximately 25% structural zeros.

Finally, following [10], we generated the read count *X*_*ij*_ for samples with 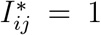 from a negative-binomial distribution with mean 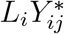 and overdispersion parameter 1, where the library size *L*_*i*_ was sampled from *N* (10, 000, 10, 000^2^) and left-truncated at 5,000. We set *X*_*ij*_ = 0 for samples with 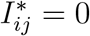. We set the number of taxa to *J* = 100 for these simulations. Across all simulation settings, we evaluated TestNet-PA against Fisher’s exact test for detecting co-occurrence dependence. Prior to analysis, we used prevalence-based filtering to remove taxa with too few observed presences or absences. Because sampling zeros make observed prevalence systematically lower than true prevalence, we used asymmetric thresholds, retaining taxa with observed prevalence between 5% and 80%. In the second set of simulations, we further benchmarked TestNet-PA against co-abundance methods, including TestNet, Spearman’s correlation test applied to relative abundance (Spearman-RA), SparCC [7], and the linear and nonlinear versions of SECOM [9]. For these methods, filtering was applied only at the lower end, excluding taxa with prevalence below 5%. For SparCC, which does not provide *p*-values, we classified taxon pairs with an absolute estimated correlation exceeding 0.3 as significant. For all other methods that provide *p*-values, we applied the BH procedure to determine significance.

We evaluated method performance using four metrics: the false positive rate (FPR, or type I error) at the 1% nominal level; empirical FDR at the 1% nominal level in the main simulations, and 10% in the second simulation setting to better distinguish sensitivity across the co-abundance methods; sensitivity (or recall) at the corresponding FDR level; and the area under the precision–recall curve (PRAUC, or simply AUC). For a fair comparison, sensitivity was evaluated for all methods using the same set of truly dependent taxon pairs. Results were based on 100 simulation replicates, except under the global null setting corresponding to the identity covariance structure. In this setting, the FDR for each replicate can take only the values 0 or 1 and is therefore inherently unstable; we thus increased the number of replicates to 1,000 to obtain more stable estimates.

### Simulation results

#### TestNet-PA versus Fisher’s exact test

Figures 2–4 summarize the results from the main simulation settings corresponding to the three templates. Across all scenarios, TestNet-PA consistently controlled the FPR and FDR below their nominal levels, with both error rates approaching the nominal levels as sample size increased. It also achieved higher AUC in most scenarios. In the gut- and vaginal-based simulations, TestNet-PA attained sensitivity comparable to Fisher’s test, despite Fisher’s increasing anti-conservativeness with sample size. The contrast was more pronounced in the oral-based simulations, where strong confounding by library size led Fisher’s test to reject excessively, producing both severely inflated FDR and deceptively high sensitivity.

**Figure 2:**
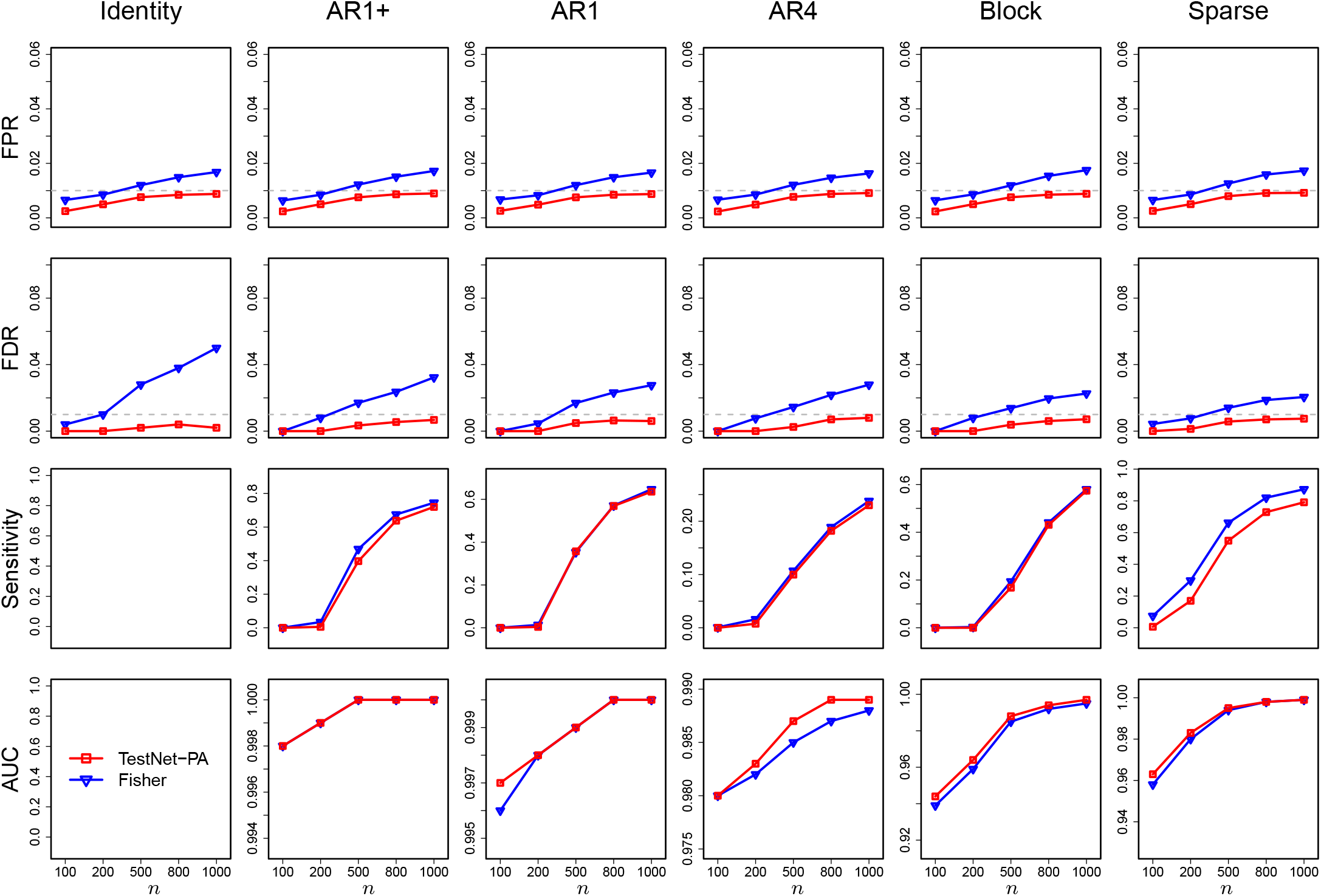
Simulation results for co-occurrence dependence inference based on the gut data template.The gray dashed line denotes the 1% nominal level for both FPR and FDR.

**Figure 3:**
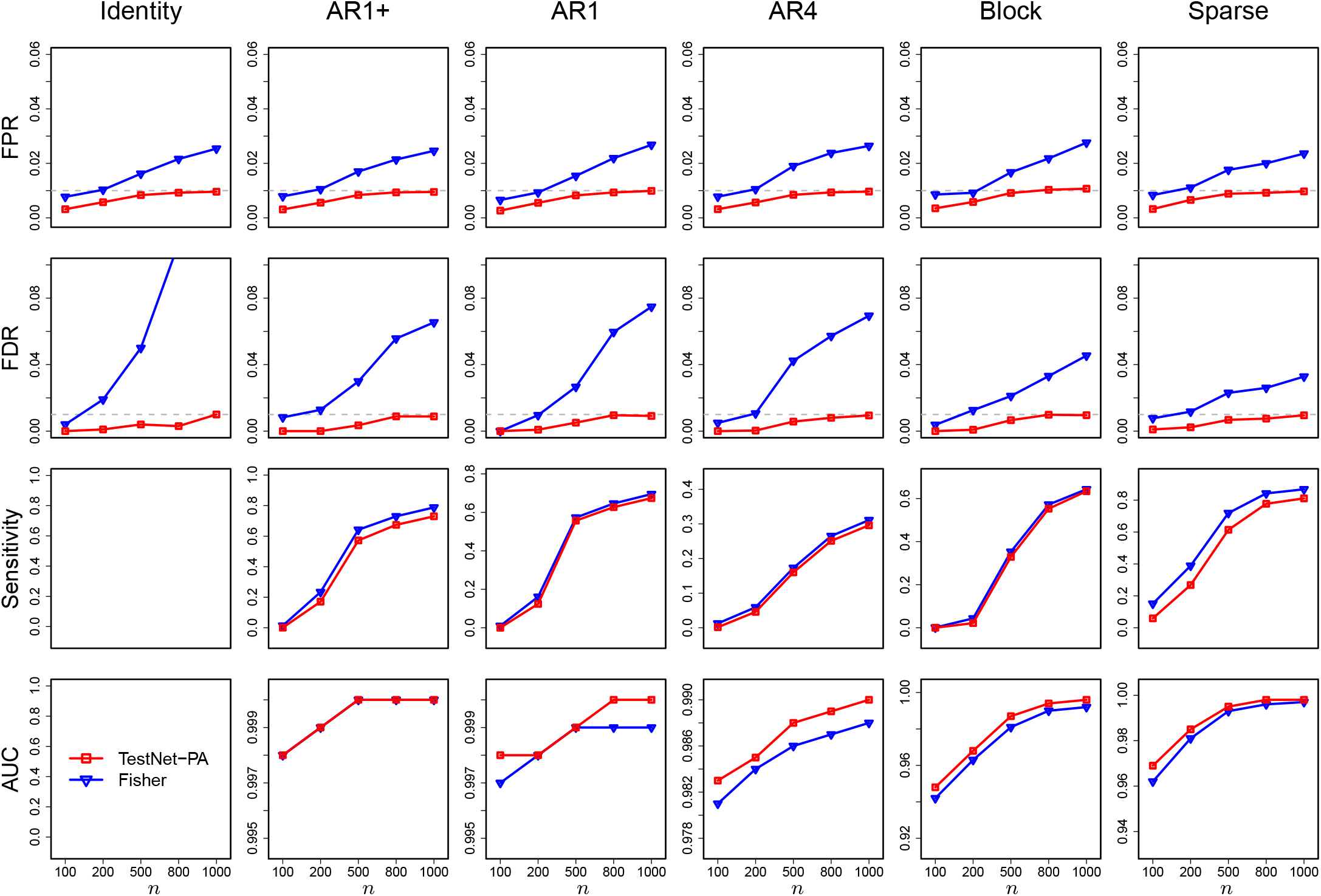
Simulation results for co-occurrence dependence inference based on the vaginal data template.The gray dashed line denotes the 1% nominal level for both FPR and FDR.

**Figure 4:**
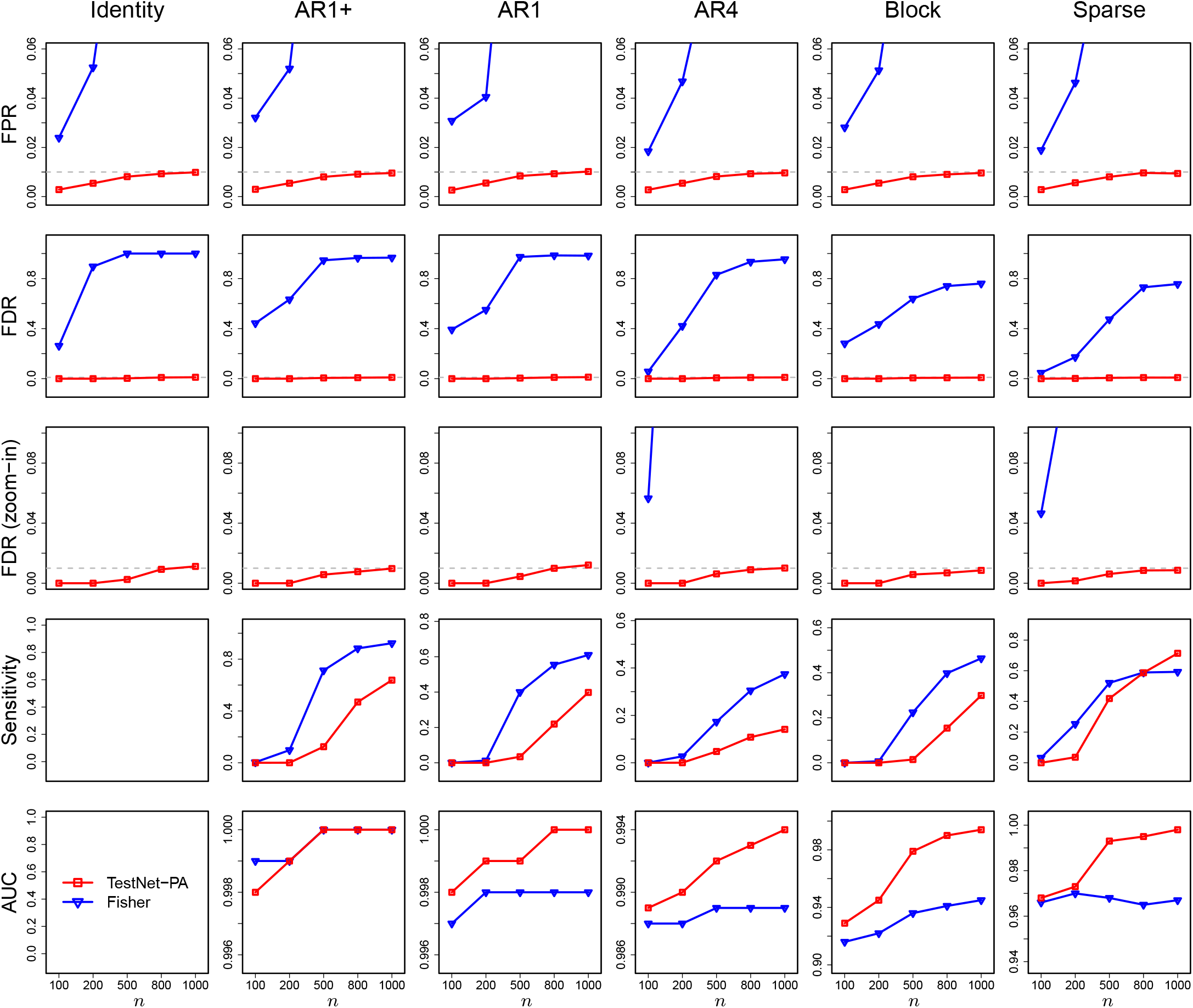
Simulation results for co-occurrence dependence inference based on the oral data template. The gray dashed line denotes the 1% nominal level for both FPR and FDR.

Figure 5 compares the dependent taxon pairs, together with their directions of dependence, identified by TestNet-PA and Fisher’s exact test in a representative replicate for each covariance structure containing both positive and negative dependencies. Strikingly, pairs uniquely detected by Fisher’s test were predominantly positive, whereas those uniquely detected by TestNet-PA were mostly negative. This pattern suggests that Fisher’s test tends to produce excess positive dependence, largely driven by library size confounding.

**Figure 5:**
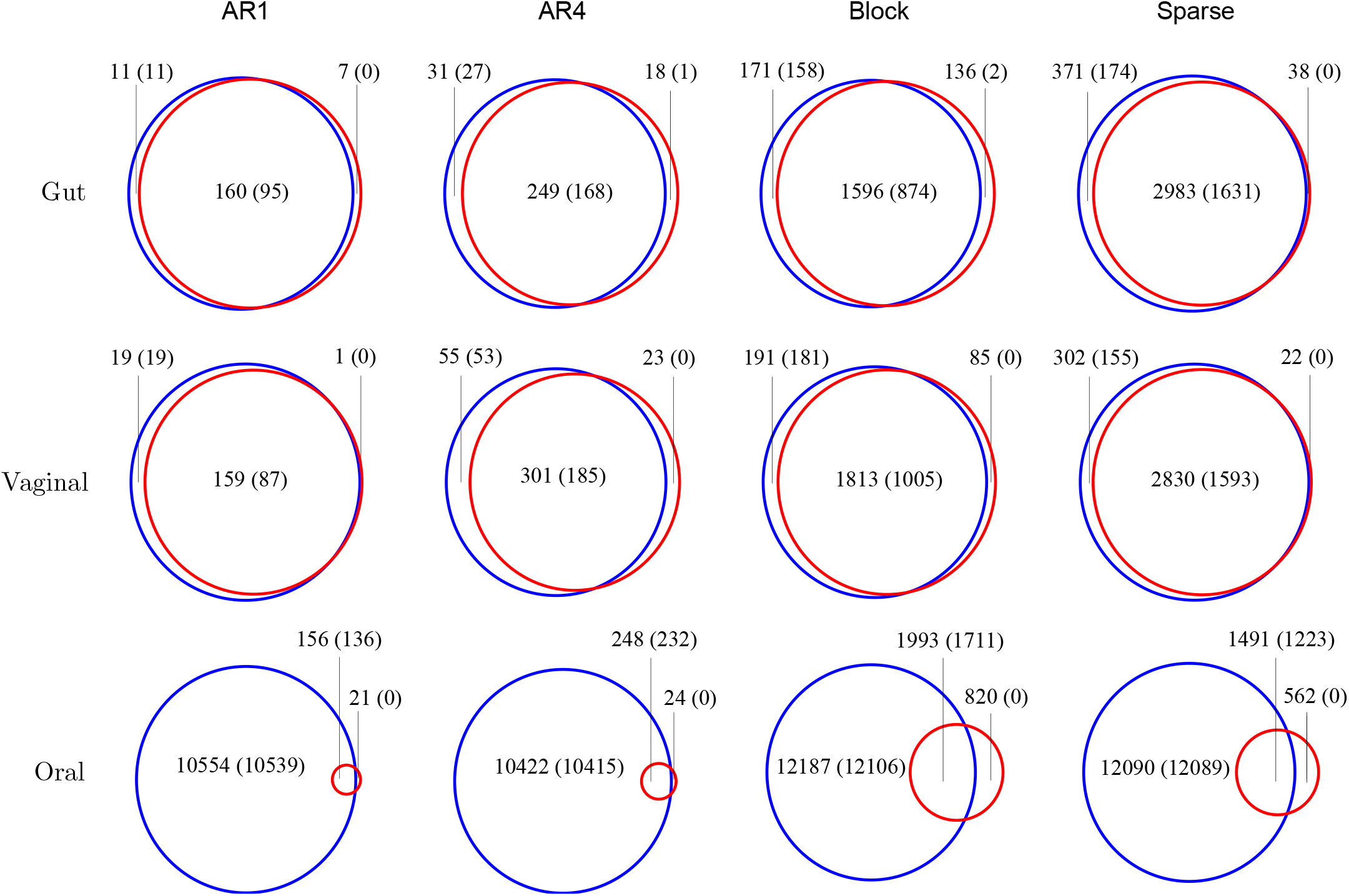
Co-occurrence dependencies identified by TestNet-PA (red circle) and Fisher’s exact test (blue circle), with the numbers of positive pairs shown in parentheses. Results are based on a representative simulation replicate with sample size *n* = 1, 000 under each covariance structure. The AR1+ setting is omitted because all of its dependencies are positive.

#### TestNet-PA versus TestNet and other co-abundance methods

Figure 6 further highlights the distinct value of TestNet-PA compared with co-abundance methods, including TestNet. While TestNet retained some ability to detect co-occurrence dependence, TestNet-PA was substantially more sensitive to this type of signal. As expected, Spearman-RA showed inflated FDR due to the compositional constraint inherent in relative abundance data. SparCC and SECOM-nonlinear exhibited no power, whereas SECOM-linear suffered from severe error-rate inflation. Fisher’s test again became increasingly anti-conservative with increasing sample size.

**Figure 6:**
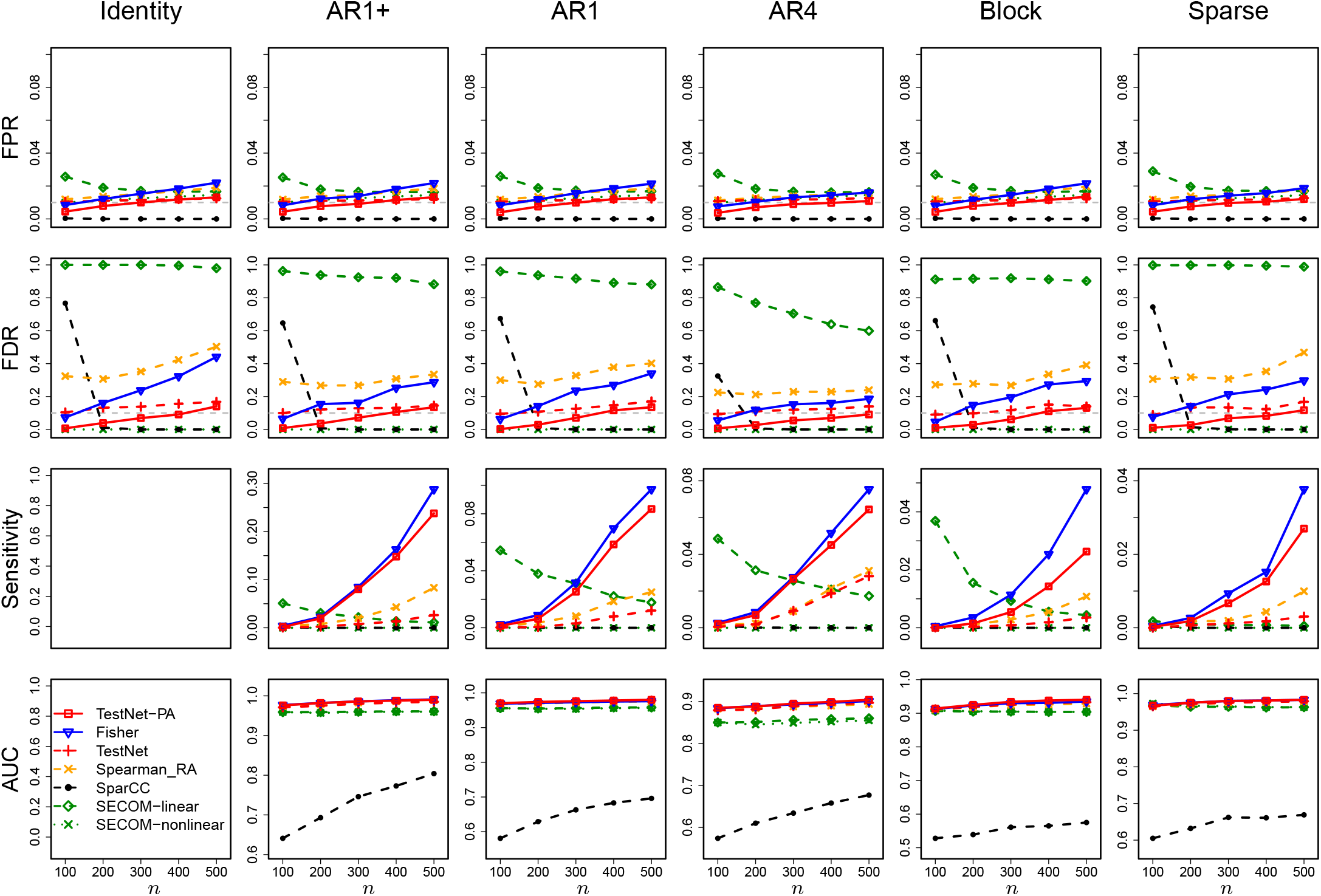
Simulation results comparing co-occurrence and co-abundance dependence inference under the second set of simulation settings. Solid lines denote co-occurrence methods, while dashed and dotted lines denote co-abundance methods. The gray dashed line marks the nominal thresholds of 1% for FPR and 10% for FDR.

#### *β* and standard error estimation

Beyond the FDR and sensitivity results across all taxon pairs, we further examined taxon-pair-specific estimation of *β* to better understand the differences between TestNet-PA and Fisher’s exact test. Using the oral data template under the identity covariance structure, for which all taxon pairs were null, we selected six representative pairs spanning the prevalence combinations (low, low), (median, median), (high, high), (low, median), (low, high), and (median, high). Here, low, median, and high prevalence correspond to approximately 0.3, 0.5, and 0.8, respectively. We also varied the sample size from 100 to 1,000. Figures 7(a) and S2(a) display the distributions, across 1,000 simulation replicates, of three estimators of *β*: the true-status-based estimator, defined as the log odds ratio computed from the true presence–absence statuses; the observed-status-based estimator, defined as the log-odds ratio computed from the observed presence–absence statuses; and the TestNet-PA estimator. As expected under the null, both the true-status-based and TestNet-PA estimates were centered at zero. By contrast, the observed-status-based estimates became increasingly positively biased as taxon prevalence or sample size increased, illustrating the bias underlying Fisher’s test.

**Figure 7:**
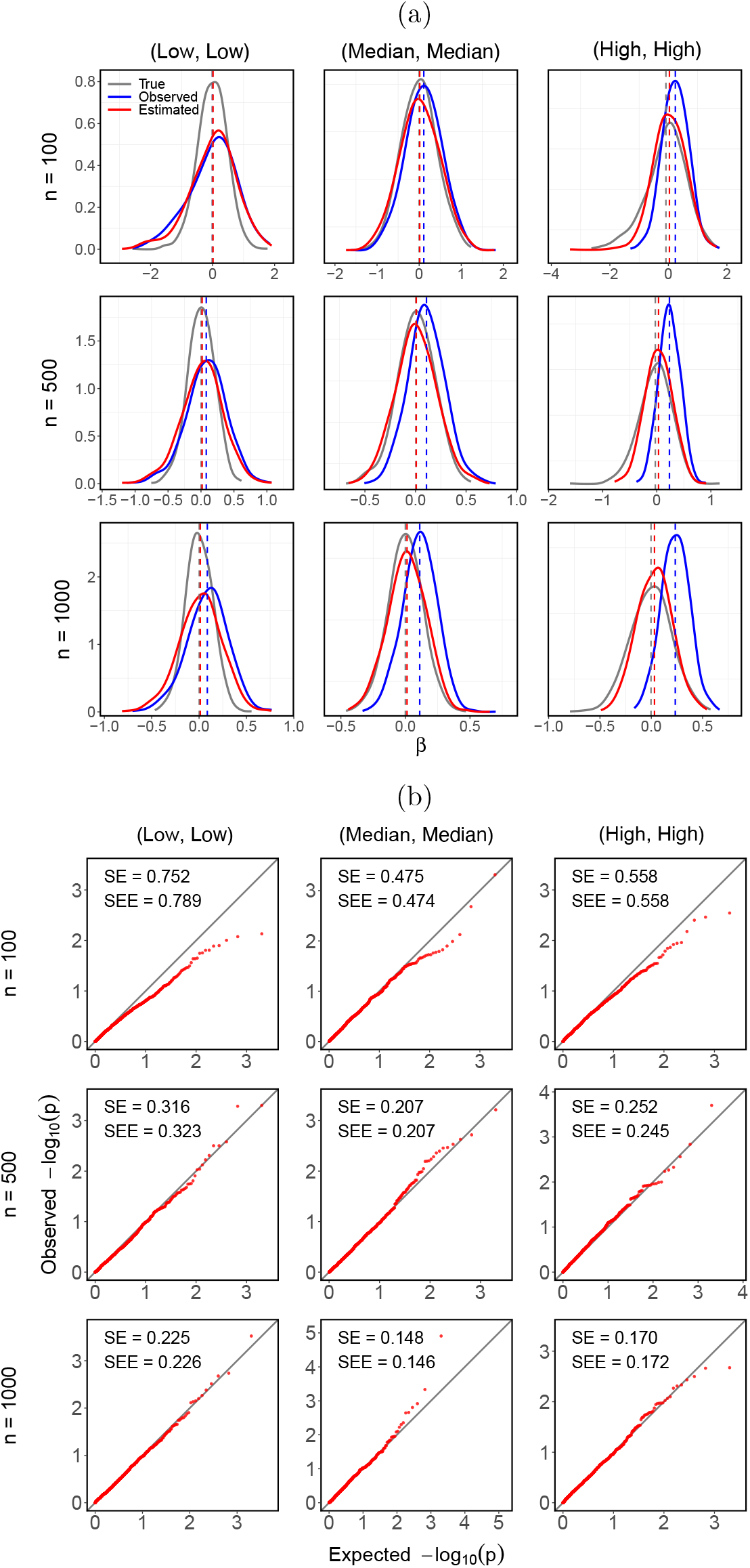
(a) Distributions of the true-status-based, observed-status-based, and TestNet-PA estimates of *β* across 1,000 simulation replicates based on the oral data template and identity covariance structure, for the three taxon pairs comprising taxa with similar prevalence levels. Each dashed line marks the mean of *β* estimates shown in the corresponding color. (b) Quantile-quantile (QQ) plots of TestNet-PA *p*-values against the reference Unif[0, 1] distribution (indicated by the gray line). SE denotes the empirical standard deviation of 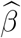 across 1,000 replicates, and SEE denotes the mean standard error estimate of 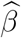.

Nonetheless, small discrepancies remained between the means of the true-status-based and TestNet-PA estimates of *β*, particularly for high-prevalence pairs. We hypothesized that these discrepancies arose from upward bias introduced during imputation. To assess this, we constructed an oracle TestNet-PA estimator of *β* by replacing the imputed values in TestNet-PA with the means of the potential relative abundances. The oracle estimator largely eliminated the remaining discrepancies (Figure S3).

We also examined the accuracy of the standard error estimates produced by TestNet-PA. As shown in Figures 7(b) and S2(b), the mean standard error estimates tended to be slightly larger than the empirical standard deviation of *β* for low-prevalence pairs and small sample sizes, but became closely aligned with it as taxon prevalence or sample size increased. This pattern supports the validity of our standard error estimation. The corresponding quantile–quantile (QQ) plots of the TestNet-PA *p*-values showed a consistent pattern, with the *p*-values being either slightly conservative or well calibrated against the reference distribution.

### Real data analysis

Before analyzing the three real datasets, we filtered out taxa with prevalence below 5% or above 80%, leaving 277, 201, and 384 taxa, respectively (Table 1). We then applied TestNet-PA and Fisher’s exact test to infer co-occurrence networks, with the nominal FDR level set at 1%. As an initial diagnostic, we inspected the *p*-value distributions. Figure S4 shows that the TestNet-PA *p*-values exhibited the expected distributional patterns, whereas Fisher’s test produced an excess of *p*-values near 1 in all three datasets.

Table 1 and Figure 8 summarize the co-occurrence dependencies detected by TestNet-PA and Fisher’s exact test. In the IBDMDB dataset, which had a relatively small sample size, TestNet-PA detected only about half as many pairs as Fisher’s test, suggesting that a substantial fraction of the additional pairs identified by Fisher’s test may be false positives. In the larger MOMS-PI dataset, TestNet-PA recovered a much larger set of pairs, nearly all of which overlapped with those identified by Fisher’s test. The contrast was most striking in the ORIGINS dataset, where the sample size was limited and library-size confounding was pronounced (Figure 1(b)). TestNet-PA detected relatively few pairs after accounting for this confounding, whereas Fisher’s test produced a very large number of detections. These patterns are consistent with the performance trends observed in our simulations, where TestNet-PA consistently controlled error rates, had moderate sensitivity at sample sizes below 500, and gained sensitivity rapidly with increasing sample size.

**Figure 8:**
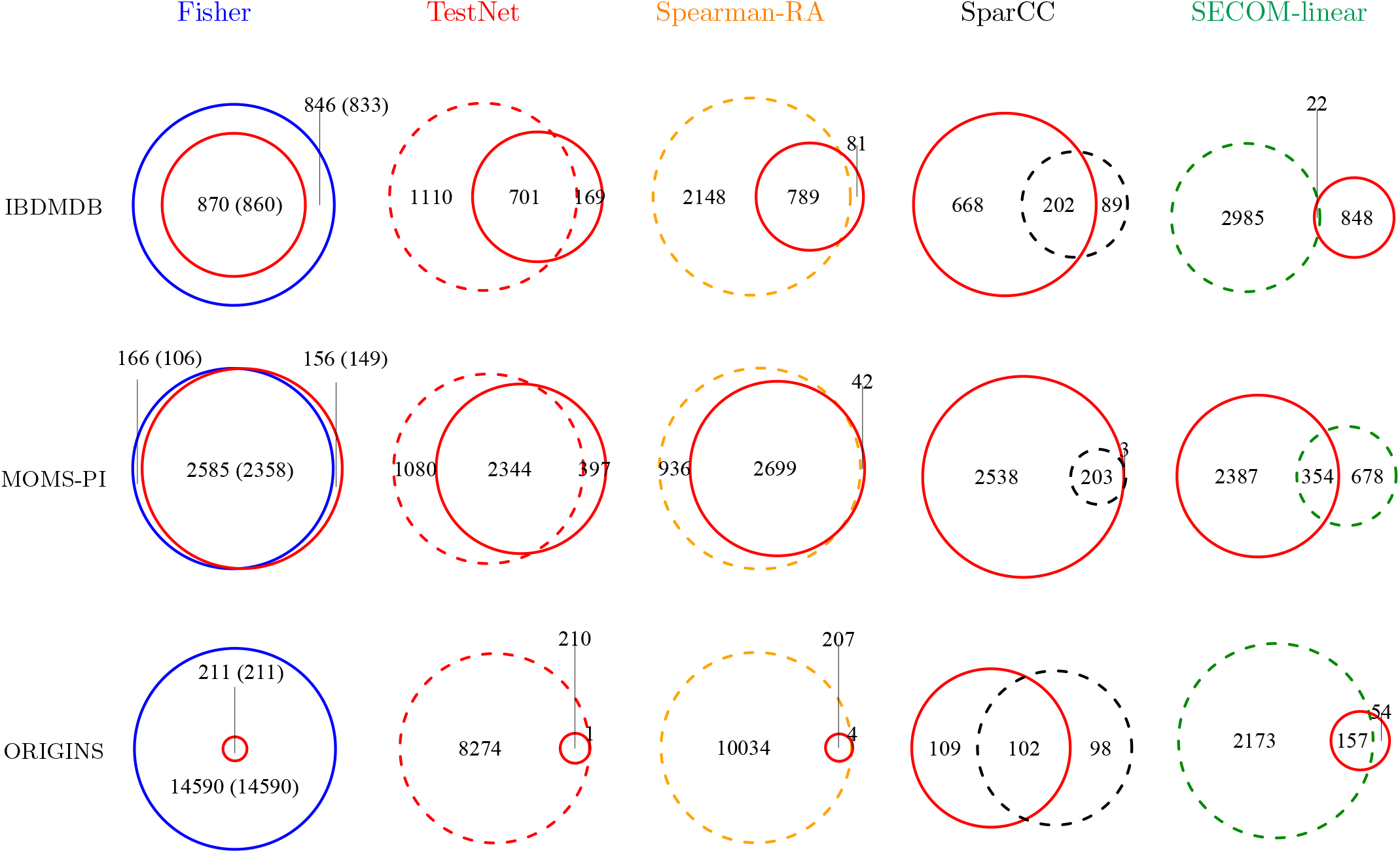
Co-occurrence and co-abundance dependencies identified in the real datasets by TestNet-PA and the comparison methods. TestNet-PA is consistently represented by the solid red circle, while the other circle denotes the method being compared; numbers in parentheses indicate positive dependencies. Co-occurrence and co-abundance methods are represented by solid and dashed circles, respectively. SECOM-nonlinear yielded no detections for any dataset and is therefore omitted.

Table 1 and Figure 8 further reveal a strong predominance of positive dependencies across all three datasets. Two factors may contribute to this pattern. First, positive dependencies may genuinely be more common in microbial communities. Consistent with this interpretation, even the pairs uniquely detected by TestNet-PA were predominantly positive, a pattern opposite to that observed in our simulations. Second, negative dependencies appear inherently more difficult to detect under library-size confounding. As shown in Figure S5, TestNet-PA consistently had lower sensitivity for negative than for positive dependencies across all simulation settings. This imbalance was most pronounced for the oral data template, where library size confounding was strongest.

Importantly, real microbiome data may contain both co-occurrence and co-abundance dependence, and the corresponding methods may capture complementary signals. With sufficient sample size, however, the two may not be empirically distinguishable, because either may produce signals detectable by both classes of methods. We therefore also applied Test-Net, Spearman-RA, SparCC, SECOM-linear, and SECOM-nonlinear to infer co-abundance networks in the same datasets, with the results also summarized in Table 1 and Figure 8. To facilitate a fair comparison with the co-occurrence analysis, we restricted all methods to the common set of taxa retained after co-occurrence filtering, since co-abundance methods may additionally include high-prevalence taxa. An initial diagnostic revealed well-behaved *p*-value distributions for TestNet and Spearman-RA, in contrast to SECOM-linear and SECOM-nonlinear, which exhibited an excess of *p*-values near 1 across all three datasets (Figure S4).

Figure 8 indeed shows that TestNet identified many pairs not detected by TestNet-PA, but it also missed a modest number of pairs uniquely identified by TestNet-PA. The overlap between the two methods was greatest in the MOMS-PI dataset, which had a substantially larger sample size than the other two datasets. In the ORIGINS dataset, by contrast, TestNet-PA discoveries constituted only a small subset of those identified by TestNet, possibly because strong library-size confounding substantially reduced the power of TestNet-PA. The remaining co-abundance methods showed patterns broadly consistent with their simulation performance, either through inflated error rates or limited power: Spearman-RA generated the largest or among the largest numbers of discoveries across the datasets; SECOM-linear showed almost no overlap with TestNet-PA in IBDMDB and only limited overlap in MOMS-PI; SparCC identified very few pairs; and SECOM-nonlinear identified none.

Figure 9 displays the co-occurrence networks inferred by TestNet-PA from the three datasets. Because taxonomic annotations were unavailable for IBDMDB and MOMS-PI, we restricted biological interpretation to the ORIGINS network. One notable module connected *Rhodoplanes, Marinifilum*, and *Oceanibium*, three genera with a common aquatic or sediment-associated ecological signature. *Marinifilum* and *Oceanibium* are primarily linked to marine and coastal environments, whereas *Rhodoplanes* also occurs in brackish and waterlogged habitats [38, 39]. Given that these genera are primarily characterized from environmental rather than oral habitats, their co-occurrence may reflect shared aquatic-associated exposure rather than direct microbial interactions. A second module comprised *Aquidulcibacter, Rhodomicrobium*, and *Frankia* and spanned a broader freshwater–soil–plant ecological gradient. *Aquidulcibacter* is associated with freshwater cyanobacterial aggregates [40], *Frankia* is a characteristic soil and rhizosphere bacterium with nitrogen-fixing plant symbioses [41], and *Rhodomicrobium* bridges these ecological niches through its occurrence in freshwater soils and sediments and plant-associated environments [42]. This module may therefore capture a common environmental-exposure signature arising from freshwater-, soil-, or plant-associated microbial sources.

**Figure 9:**
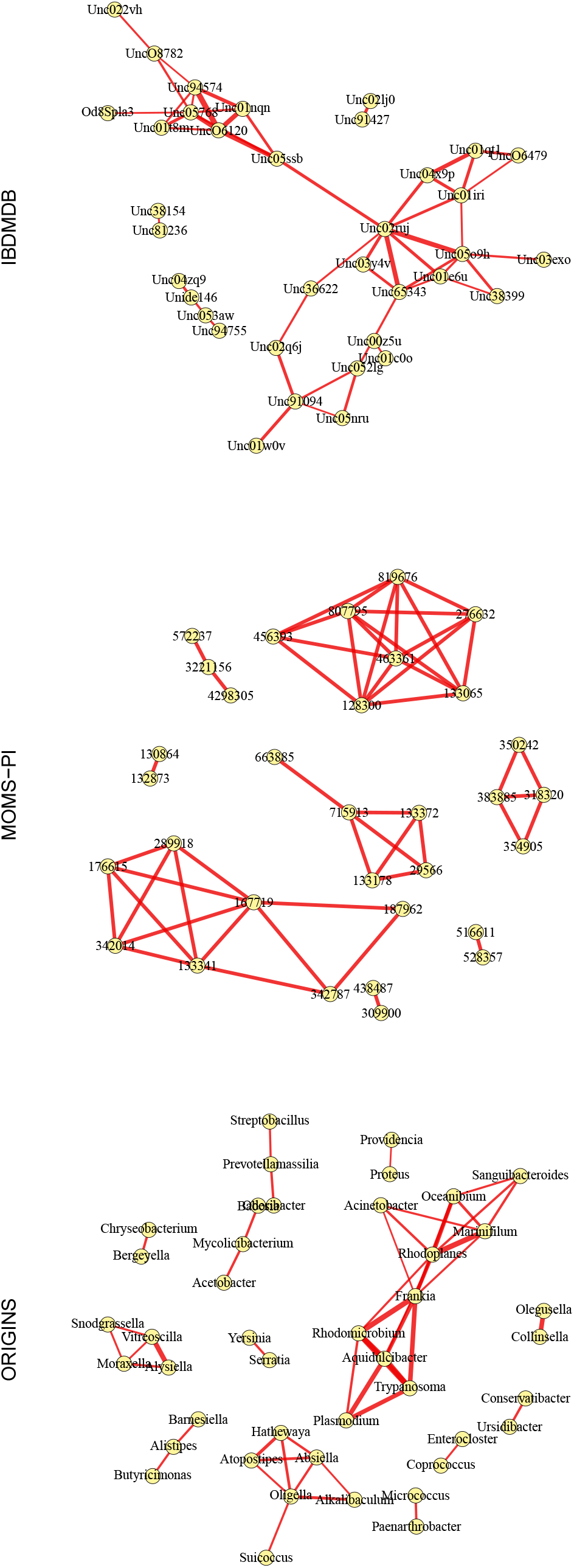
Microbial co-occurrence networks inferred by TestNet-PA, constructed using the 50 most significant taxon pairs in each real dataset. All displayed edges correspond to positive dependencies, with thicker edges indicating greater statistical significance, quantified by − log_10_(*q*-value).

## Discussion

We presented TestNet-PA, a novel method for testing pairwise presence–absence dependence among microbial taxa. Starting directly from read count data, TestNet-PA accounts for library-size confounding and explicitly models the discrepancy between observed detection statuses and the underlying true presence–absence states. Presence–absence analysis provides a valuable and robust alternative to abundance-based approaches for microbiome data, yet dedicated methods in this area remain scarce. TestNet-PA addresses this gap and, to our knowledge, is the first statistically principled method designed specifically for microbial network inference based on presence–absence dependence. Across simulations, TestNet-PA controlled false discoveries while retaining good sensitivity, whereas Fisher’s exact test became increasingly anti-conservative, particularly under strong library-size confounding; the real-data analyses showed corresponding differences in the inferred networks.

Co-occurrence and co-abundance networks capture different dimensions of microbial dependence and can therefore offer complementary biological insights. Presence–absence analysis may be relatively advantageous compared with abundance-based methods for sparse taxa, whereas abundance-based methods may be more powerful for taxa with substantial quanti-tative variation. These considerations motivate future development of omnibus or integrative approaches that combine both sources of information.

Beyond microbiome applications, TestNet-PA may have broader utility in sequencing-based studies where presence–absence patterns are biologically informative. In single-cell RNA-seq data, for example, it could identify gene pairs that tend to be jointly ON or OFF across cells, potentially capturing shared cell-type specificity, coordinated regulatory programs, or common activation states. TestNet-PA may also extend naturally to TCR/BCR sequencing, where co-occurrence among clonotypes could help characterize shared antigen specificity, convergent immune responses, and disease-associated repertoire signatures.

TestNet-PA is implemented in our R package TestNet [10], available on CRAN and GitHub at https://github.com/yijuanhu/TestNet. The method is computationally efficient, requiring only 16.4, 14.7, and 50.0 seconds to analyze the IBDMDB, MOMS-PI, and ORIGINS datasets, respectively, on a MacBook Pro equipped with an Apple M2 chip and 16 GB of memory.

TestNet-PA currently targets marginal pairwise dependence, so detected edges may represent indirect associations induced by shared host, environmental, or technical factors rather than direct microbial interactions. In addition, inference relies on a working model for sampling-zero probabilities and on imputation of unobserved relative abundances; although our simulations suggest robustness to these approximations, substantial departures from the underlying assumptions may affect calibration. Extensions to covariate-adjusted, clustered, and conditional-dependence settings represent important directions for future work. Together, these developments could further broaden the utility of presence–absence-based network inference across microbiome and other sequencing studies.

