## Supplementary material for "TestNet-PA: Sampling-Zero-Aware Inference of Microbial Presence-Absence Networks"

### Supplementary Materials

#### Attenuation of dependence parameters from omitted random effects

Let $U_{ik} = L_i(\overline{Y}_k - Y^*_{ik})$ denote the random effect induced by imputation, or simply the imputation error, and assume that it has mean of zero: $\mathrm{E}(U_{ik} \mid L_i) = 0$, where $L_i$ denotes the library size. It is also reasonable to assume that $U_{ik} \perp I^*_{ij} \mid I^*_{ik}, L_i$. The true and working sampling-zero probabilities are related by

$$q_{ik} = \exp(-L_i Y^*_{ik}) = \exp[-L_i\overline{Y}_k + L_i(\overline{Y}_k - Y^*_{ik})] = \widehat{q}_{ik}\exp(U_{ik}).$$

Because $U_{ik}$ is unobserved and omitted from the working model, integrating over its distribution yields $\mathrm{E}(q_{ik} \mid L_i) = \widehat{q}_{ik}\mathrm{E}(\exp(U_{ik}) \mid L_i)$. By Jensen's inequality, $\mathrm{E}[\exp(U_{ik}) \mid L_i] \geq \exp[\mathrm{E}(U_{ik} \mid L_i)] = 1$, and hence

$$\mathrm{E}(q_{ik} \mid L_i) \geq \widehat{q}_{ik}. \tag{S1}$$

The key device in the proof is an auxiliary thinning step that links the true and working detection mechanisms. Define the pseudo-true status $I^\dagger_{ik}$ by thinning the true status $I^*_{ik}$ with retention probability

$$P(I^\dagger_{ik} = 1 \mid I^*_{ik} = 1, L_i) = \lambda_{ik} \equiv \frac{1 - \mathrm{E}(q_{ik} \mid L_i)}{1 - \widehat{q}_{ik}},$$

where (S1) ensures that $0 \leq \lambda_{ik} \leq 1$. Applying the working detection with probability

$$P(I_{ik} = 1 | I^\dagger_{ik} = 1, L_i) = 1 - \widehat{q}_{ik} \tag{S2}$$

to the thinned status $I^\dagger_{ik}$ exactly reproduces the true detection with probability

$$P(I_{ik} = 1 | I^*_{ik} = 1, L_i) = 1 - \mathrm{E}(q_{ik} \mid L_i) = \lambda_{ik}(1 - \widehat{q}_{ik}). \tag{S3}$$

Thus, the true detection mechanism can be represented by the two-stage procedure as

$$I^*_{ik} \xrightarrow[\text{Extra thinning}]{\lambda_{ik}} I^\dagger_{ik} \xrightarrow[\text{Working detection}]{1-\widehat{q}_{ik}} I_{ik}.$$

To establish attenuation of the dependence parameter between the two latent statuses, $I^*_{ij}$ and $I^*_{ik}$, we proceed in two steps. In step 1, consider the probability of detecting taxon $k$ given the true status of taxon $j$ in sample $i$, $P(I_{ik} = 1 \mid I^*_{ij} = s, L_i)$, for $s \in \{0, 1\}$. As $I_{ik} = 1$ requires $I^*_{ik} = 1$, under the true model,

$$P(I_{ik} = 1 \mid I^*_{ij} = s, L_i) = P(I_{ik} = 1 \mid I^*_{ik} = 1, I^*_{ij} = s, L_i)P(I^*_{ik} = 1 \mid I^*_{ij} = s, L_i). \tag{S4}$$

The first factor equals (S3), because the true detection of taxon $k$ does not depend on the true status of the other taxon $j$. As $I_{ik} = 1$ requires $I^\dagger_{ik} = 1$, under the working model,

$$P(I_{ik} = 1 \mid I^*_{ij} = s, L_i) = P(I_{ik} = 1 \mid I^\dagger_{ik} = 1, I^*_{ij} = s, L_i)P(I^\dagger_{ik} = 1 \mid I^*_{ij} = s, L_i), \tag{S5}$$

where the first factor equals (S2), because by construction the working detection does not depend on the true status of the other taxon. Equating (S4) and (S5) therefore yields

$$P(I^\dagger_{ik} = 1 \mid I^*_{ij} = s, L_i) = \lambda_{ik} P(I^*_{ik} = 1 \mid I^*_{ij} = s, L_i). \tag{S6}$$

We next introduce logistic models involving the parameters of interest. Assume that the true co-occurrence model does not vary with $L_i$: $P(I^*_{ik} = 1 \mid I^*_{ij} = s, L_i) = P(I^*_{ik} = 1 \mid I^*_{ij} = s) = \text{expit}(\alpha + \beta s)$, where the last equality follows from (1), and $\text{expit}(\eta) = \exp(\eta)/[1+\exp(\eta)]$. Marginalize (S6) over $L_i$ gives

$$P(I^\dagger_{ik} = 1 \mid I^*_{ij} = s) = \text{E}(\lambda_{ik} \mid I^*_{ij} = s)\, \text{expit}(\alpha + \beta s). \tag{S7}$$

We further assume that $L_i$ is independent of $I^*_{ij}$. Since $\lambda_{ik}$ is a function of $L_i$, its conditional mean does not depend on $s$; denote this common value by $\overline{\lambda}_k = \text{E}(\lambda_{ik} \mid I^*_{ij} = s)$. For the left-hand side of (S7), because $I^\dagger_{ik}$ and $I^*_{ij}$ are both binary, their conditional relationship can be represented exactly by a two-parameter logistic model: $P(I^\dagger_{ik} = 1 \mid I^*_{ij} = s) = \text{expit}(\alpha_k + \beta_k s)$. Thus, (S7) becomes

$$\text{expit}(\alpha_k + \beta_k s) = \overline{\lambda}_k\, \text{expit}(\alpha + \beta s). \tag{S8}$$

We then establish attenuation of $\beta_k$ relative to $\beta$. From (S8),

$$\beta_k = \text{logit}[\overline{\lambda}_k\, \text{expit}(\alpha + \beta)] - \text{logit}[\overline{\lambda}_k\, \text{expit}(\alpha)].$$

Define $h_k(\eta) = \text{logit}[\overline{\lambda}_k \,\text{expit}(\eta)]$, so that $\beta_k = h_k(\alpha+\beta) - h_k(\alpha)$. By the mean-value theorem, for some $\xi$ between $\alpha$ and $\alpha+\beta$,

$$\beta_k = h_k'(\xi)\beta.$$

Writing $p = \text{expit}(\eta)$, we have

$$h_k'(\eta) = \frac{d}{d\eta}\left[\log(\overline{\lambda}_k p) - \log(1-\overline{\lambda}_k p)\right] = \frac{1}{p}\frac{dp}{d\eta} + \frac{\overline{\lambda}_k}{1-\overline{\lambda}_k p}\frac{dp}{d\eta} = \frac{1-p}{1-\overline{\lambda}_k p} \in (0,1].$$

Thus, $\beta_k$ is attenuated toward zero relative to $\beta$.

In step 2, consider $P(I_{ij}=1|I_{ik}^\dagger = s, L_i)$, for $s \in \{0,1\}$, in a similar manner. First, define $I_{ij}^\dagger$ and $\lambda_{ij}$ analogously for taxon $j$. Under the true model, $P(I_{ij}=1|I_{ik}^\dagger = s, L_i) = P(I_{ij} = 1|I_{ij}^* = 1, I_{ik}^\dagger = s, L_i)P(I_{ij}^* = 1|I_{ik}^\dagger = s, L_i)$. The first factor equals the taxon-$j$ analogue of (S3), because the true detection of taxon $j$ does not depend on the pseudo-true status of the other taxon $k$. Under the working model, $P(I_{ij}=1|I_{ik}^\dagger = s, L_i) = P(I_{ij} = 1|I_{ij}^\dagger = 1, I_{ik}^\dagger = s, L_i)P(I_{ij}^\dagger = 1|I_{ik}^\dagger = s, L_i)$, where the first factor equals the taxon-$j$ analogue of (S2), because by construction the working detection does not depend on the pseudo-true status of the other taxon. Equating the two models yields

$$P(I_{ij}^\dagger = 1 \mid I_{ik}^\dagger = s, L_i) = \lambda_{ij} P(I_{ij}^* = 1 \mid I_{ik}^\dagger = s, L_i).$$

We next introduce logistic models involving the parameters of interest. By iterated expectation, the marginal relationship can be written as

$$P(I_{ij}^\dagger = 1 \mid I_{ik}^\dagger = s) = \text{E}\left[P(I_{ij}^\dagger = 1 \mid I_{ik}^\dagger = s, L_i) \mid I_{ik}^\dagger = s\right] = \text{E}\left[\lambda_{ij} P(I_{ij}^* = 1 \mid I_{ik}^\dagger = s, L_i) \mid I_{ik}^\dagger = s\right]$$
$$= \text{E}\left[\lambda_{ij}\text{E}(I_{ij}^* \mid I_{ik}^\dagger = s, L_i) \mid I_{ik}^\dagger = s\right] = \text{E}(\lambda_{ij} I_{ij}^* \mid I_{ik}^\dagger = s) = P(I_{ij}^* = 1 \mid I_{ik}^\dagger = s)\text{E}(\lambda_{ij} \mid I_{ij}^* = 1, I_{ik}^\dagger = s).$$

Assume that thinning of taxon $j$ is nondifferential with respect to the pseudo-true status of taxon $k$, so that $\text{E}(\lambda_{ij} \mid I_{ij}^* = 1, I_{ik}^\dagger = s) = \overline{\lambda}_j$ does not depend on $s$. Then

$$P(I_{ij}^\dagger = 1 \mid I_{ik}^\dagger = s) = \overline{\lambda}_j P(I_{ij}^* = 1 \mid I_{ik}^\dagger = s).$$

From the first step, the log odds ratio between $I_{ij}^*$ and $I_{ik}^\dagger$ is $\beta_k$. By symmetry of the odds ratio, we can write

$$P(I_{ij}^* = 1 \mid I_{ik}^\dagger = s) = \text{expit}(\alpha_j + \beta_k s).$$

Using (4) and the symmetry of the odds ratio, the quasi-likelihood parameter $\widetilde{\beta}$ satisfies

$$P(I_{ij}^{\dagger} = 1 \mid I_{ik}^{\dagger} = s) = \text{expit}(\widetilde{\alpha}_j + \widetilde{\beta} s)$$

for some intercept $\widetilde{\alpha}_j$. Hence,

$$\text{expit}(\widetilde{\alpha}_j + \widetilde{\beta} s) = \overline{\lambda}_j \, \text{expit}(\alpha_j + \beta_k s).$$

Applying the same argument as in step 1 gives that $\widetilde{\beta}$ is attenuated toward zero relative to $\beta_k$. Combining the two steps, the quasi-likelihood log odds ratio $\widetilde{\beta}$ is attenuated toward zero relative to the true log odds ratio $\beta$.

#### EM algorithm

At the $t$-th iteration, let $\widetilde{\Theta}^{(t)}$ denote the current parameter estimate. In the E-step, we calculate the posterior probabilities of the latent true presence–absence status $I_i^*$ given the observed status $I_i$ and $\widetilde{\Theta}^{(t)}$. When $I_i = 11$, the latent true status is known with certainty, so that $P(I_i^* = 11 \mid I_i = 11) = 1$. When $I_i = 10$, however, the latent true status may be either '11' or '10'. Under the working model, the corresponding probabilities are

$$\begin{aligned} p_{i,11|10}^{(t)} \equiv P\left(I_i^* = 11 \mid I_i = 10, \widetilde{\Theta}^{(t)}\right) &= \frac{P\left(I_i = 10 \mid I_i^* = 11\right) \widetilde{\theta}_{11}^{(t)}}{P(I_i = 10)} \\ &= \frac{(1 - \widehat{q}_{ij})\, \widehat{q}_{ik}\, \widetilde{\theta}_{11}^{(t)}}{(1 - \widehat{q}_{ij})\, \widehat{q}_{ik}\, \widetilde{\theta}_{11}^{(t)} + (1 - \widehat{q}_{ij})\, \widetilde{\theta}_{10}^{(t)}} = \frac{\widehat{q}_{ik}\, \widetilde{\theta}_{11}^{(t)}}{\widehat{q}_{ik}\, \widetilde{\theta}_{11}^{(t)} + \widetilde{\theta}_{10}^{(t)}} \end{aligned}$$

and

$$p_{i,10|10}^{(t)} \equiv 1 - p_{i,11|10}^{(t)}.$$

When $I_i = 01$, the probabilities of $I_i^* = 11$ and $I_i^* = 01$ are

$$p_{i,11|01}^{(t)} \equiv \frac{\widehat{q}_{ij}\, \widetilde{\theta}_{11}^{(t)}}{\widehat{q}_{ij}\, \widetilde{\theta}_{11}^{(t)} + \widetilde{\theta}_{01}^{(t)}} \quad \text{and} \quad p_{i,01|01}^{(t)} \equiv 1 - p_{i,11|01}^{(t)}.$$

Finally, when $I_i = 00$, the latent true status may take any of the four possible values, with corresponding probabilities

$$p_{i,11|00}^{(t)} \equiv c_i^{-1} \widehat{q}_{ij}\, \widehat{q}_{ik}\, \widetilde{\theta}_{11}^{(t)}, \quad p_{i,10|00}^{(t)} \equiv c_i^{-1} \widehat{q}_{ij}\, \widetilde{\theta}_{10}^{(t)}, \quad p_{i,01|00}^{(t)} \equiv c_i^{-1} \widehat{q}_{ik}\, \widetilde{\theta}_{01}^{(t)}, \quad \text{and} \quad p_{i,00|00}^{(t)} \equiv c_i^{-1} \widetilde{\theta}_{00}^{(t)},$$

where $c_i = \widehat{q}_{ij}\, \widehat{q}_{ik}\, \widetilde{\theta}^{(t)}_{11} + \widehat{q}_{ij}\, \widetilde{\theta}^{(t)}_{10} + \widehat{q}_{ik}\, \widetilde{\theta}^{(t)}_{01} + \widetilde{\theta}^{(t)}_{00}$.

In the M-step, we maximize the $Q$-function, which is given by

$$\begin{aligned}
Q(\widetilde{\Theta} \mid \widetilde{\Theta}^{(t)}) = \sum_{i=1}^{n} \Big\{ & \mathbb{I}(I_i = 11)\, \log\widetilde{\theta}_{11} + \mathbb{I}(I_i = 10) \left[ p^{(t)}_{i,11|10}\, \log\widetilde{\theta}_{11} + p^{(t)}_{i,10|10}\, \log\widetilde{\theta}_{10} \right] \\
& + \mathbb{I}(I_i = 01) \left[ p^{(t)}_{i,11|01}\, \log\widetilde{\theta}_{11} + p^{(t)}_{i,01|01}\, \log\widetilde{\theta}_{01} \right] \\
& + \mathbb{I}(I_i = 00) \left[ p^{(t)}_{i,11|00}\, \log\widetilde{\theta}_{11} + p^{(t)}_{i,10|00}\, \log\widetilde{\theta}_{10} + p^{(t)}_{i,01|00}\, \log\widetilde{\theta}_{01} + p^{(t)}_{i,00|00}\, \log\widetilde{\theta}_{00} \right] \Big\}.
\end{aligned}$$

Rearranging the $Q$-function in terms of $\log\widetilde{\theta}$, let $T^{(t)}$ denote the corresponding coefficients. For example, the coefficient for $\log\widetilde{\theta}_{11}$ is

$$T^{(t)}_{11} = \sum_{i=1}^{n} \left[ \mathbb{I}(I_i = 11) + \mathbb{I}(I_i = 10)\, p^{(t)}_{i,11|10} + \mathbb{I}(I_i = 01)\, p^{(t)}_{i,11|01} + \mathbb{I}(I_i = 00)\, p^{(t)}_{i,11|00} \right].$$

Note that $T^{(t)}_{11} + T^{(t)}_{10} + T^{(t)}_{01} + T^{(t)}_{00} = n$. The Q-function can then be rewritten as

$$Q(\widetilde{\Theta} \mid \widetilde{\Theta}^{(t)}) = T^{(t)}_{11} \log\widetilde{\theta}_{11} + T^{(t)}_{10} \log\widetilde{\theta}_{10} + T^{(t)}_{01} \log\widetilde{\theta}_{01} + T^{(t)}_{00} \log\widetilde{\theta}_{00}.$$

Reparameterizing in terms of $(\widetilde{\pi}, \widetilde{\alpha}, \widetilde{\beta})$, we can further write the $Q$-function as

$$\begin{aligned}
Q(\widetilde{\pi}, \widetilde{\alpha}, \widetilde{\beta} | \widetilde{\pi}^{(t)}, \widetilde{\alpha}^{(t)}, \widetilde{\beta}^{(t)}) \;=\; & T^{(t)}_{11} \log\left\{ \widetilde{\pi}\ \mathrm{expit}(\widetilde{\alpha} + \widetilde{\beta}) \right\} + T^{(t)}_{10} \log\left\{ \widetilde{\pi}\big[1 - \mathrm{expit}(\widetilde{\alpha} + \widetilde{\beta})\big] \right\} \\
+ \; & T^{(t)}_{01} \log\left\{ (1 - \widetilde{\pi})\, \mathrm{expit}(\widetilde{\alpha}) \right\} + T^{(t)}_{00} \log\left\{ (1 - \widetilde{\pi})\big[1 - \mathrm{expit}(\widetilde{\alpha})\big] \right\}.
\end{aligned}$$

Maximization of this expression yields the parameter updates for the $(t+1)$-th iteration:

$$\widetilde{\pi}^{(t+1)} = \frac{T^{(t)}_{11} + T^{(t)}_{10}}{n},$$

and

$$\widetilde{\beta}^{(t+1)} = \log\left[ \frac{T^{(t)}_{11} T^{(t)}_{00}}{T^{(t)}_{10} T^{(t)}_{01}} \right], \quad \widetilde{\alpha}^{(t+1)} = \log\left[ \frac{T^{(t)}_{01}}{T^{(t)}_{00}} \right]. \tag{S9}$$

#### Firth-like correction

We introduce a bias-reducing correction motivated by Firth's method [35], which achieves first-order bias reduction through an adjustment to the score function and, in likelihood-based models, is equivalent to Jeffreys-prior penalization. In our setting, however, the observed-data quasi-likelihood has a mixture form, for which the corresponding bias-reducing adjustment does not yield a simple closed-form solution. We therefore adopt a more natural correction within the EM algorithm. Specifically, the updates in (S9) have the same form as the maximum-likelihood estimates from a standard logistic regression applied to a $2 \times 2$ table, with the $T^{(t)}$'s playing the role of cell counts. Motivated by this correspondence, at each M-step, we apply the standard Firth correction to this expected complete-data $2 \times 2$ table, replacing (S9) by the updates obtained after adding 0.5 to each cell count:

$$\widetilde{\beta}^{(t+1)} = \log\left[\frac{(T_{11}^{(t)} + 0.5)(T_{00}^{(t)} + 0.5)}{(T_{10}^{(t)} + 0.5)(T_{01}^{(t)} + 0.5)}\right], \quad \widetilde{\alpha}^{(t+1)} = \log\left[\frac{T_{01}^{(t)} + 0.5}{T_{00}^{(t)} + 0.5}\right].$$

For the marginal Bernoulli parameter $\widetilde{\pi}$, we analogously apply the Firth correction and replace $\widetilde{\pi}^{(t+1)}$ above (S9) by $\widetilde{\pi}^{(t+1)} = (T_{11}^{(t)} + T_{10}^{(t)} + 0.5)/(n+1)$.

#### Variance estimation for $\widehat{\beta}$

We show below that, under the null hypothesis ($\beta = 0$), the sandwich variance under the true data-generating model agrees with the inverse information matrix under the working quasi-likelihood to first order in the random effects.

Recall that $q_{ij} = \widehat{q}_{ij} \exp(U_{ij})$, where $U_{ij} = L_i(\overline{Y}_j - Y_{ij}^*)$, and analogously for taxon $k$. Thus, the working model corresponds to $U_{ij} = U_{ik} = 0$. Let $\mathcal{W}$ denote this working model and $\mathcal{T}$ the true model obtained after averaging over the random effects. Conditional on $L_i$, $U_{ij}$, and $U_{ik}$, the probabilities of the four observed presence–absence outcomes are smooth functions of $q_{ij}$ and $q_{ik}$, and hence of $U_{ij}$ and $U_{ik}$. For each observed outcome $r$ in $\{00, 01, 10, 11\}$, a Taylor expansion around $U_{ij} = U_{ik} = 0$ gives

$$P(I_i = r \mid L_i, U_{ij}, U_{ik}) = P_{\mathcal{W}}(I_i = r \mid L_i) + C_{ij,r}U_{ij} + C_{ik,r}U_{ik} + O\left(U_{ij}^2 + U_{ik}^2 + |U_{ij}U_{ik}|\right),$$

for some coefficients $C_{ij,r}$ and $C_{ik,r}$. Since $\mathrm{E}_{\mathcal{T}}(U_{ij} \mid L_i) = \mathrm{E}_{\mathcal{T}}(U_{ik} \mid L_i) = 0$, the linear terms vanish after averaging over the random effects. Hence,

$$P_{\mathcal{T}}(I_i = r \mid L_i) = P_{\mathcal{W}}(I_i = r \mid L_i) + O\left[\mathrm{E}_{\mathcal{T}}(U_{ij}^2 \mid L_i) + \mathrm{E}_{\mathcal{T}}(U_{ik}^2 \mid L_i)\right], \tag{S10}$$

where the cross-product term is of the same order by the Cauchy–Schwarz inequality.

Let $\widetilde{\ell}_i(\widetilde{\Theta})$ denote the contribution of sample $i$ to the quasi-likelihood in (3), and define the corresponding score vector and Hessian by

$$\widetilde{S}_i(\widetilde{\Theta}) = \frac{\partial \widetilde{\ell}_i(\widetilde{\Theta})}{\partial \widetilde{\Theta}}, \qquad \widetilde{H}_i(\widetilde{\Theta}) = \frac{\partial^2 \widetilde{\ell}_i(\widetilde{\Theta})}{\partial \widetilde{\Theta} \partial \widetilde{\Theta}^\top},$$

where $\widetilde{\Theta} = (\widetilde{\pi}, \widetilde{\alpha}, \widetilde{\beta})^\top$. We showed earlier that the null condition $\beta = 0$ corresponds to the null parameter $\widetilde{\Theta}_0 = (\widetilde{\pi}_0, \widetilde{\alpha}_0, 0)^\top$. Under the working model, $\widetilde{\ell}_i(\widetilde{\Theta})$ coincides, up to additive terms independent of $\widetilde{\Theta}$, with a proper observed-data log-likelihood. Therefore, the usual information identity holds:

$$-\mathrm{E}_{\mathcal{W}}[\widetilde{H}_i(\widetilde{\Theta}_0)] = \mathrm{E}_{\mathcal{W}}[\widetilde{S}_i(\widetilde{\Theta}_0)\widetilde{S}_i(\widetilde{\Theta}_0)^\top] \equiv \mathcal{I}_{\mathcal{W}}.$$

Now define the bread and meat of the sandwich variance under the true data-generating distribution as

$$A_{\mathcal{T}} = -\mathrm{E}_{\mathcal{T}}[\widetilde{H}_i(\widetilde{\Theta}_0)], \qquad B_{\mathcal{T}} = \mathrm{E}_{\mathcal{T}}[\widetilde{S}_i(\widetilde{\Theta}_0)\widetilde{S}_i(\widetilde{\Theta}_0)^\top].$$

Conditional on $L_i$, both $\widetilde{H}_i(\widetilde{\Theta}_0)$ and $\widetilde{S}_i(\widetilde{\Theta}_0)\widetilde{S}_i(\widetilde{\Theta}_0)^\top$ depend on the observed outcome only through the four possible values of $I_i$. Therefore, the expansion in (S10), followed by averaging over $L_i$, implies

$$A_{\mathcal{T}} = \mathcal{I}_{\mathcal{W}} + O\left[\mathrm{Var}(U_{ij}) + \mathrm{Var}(U_{ik})\right],$$

and

$$B_{\mathcal{T}} = \mathcal{I}_{\mathcal{W}} + O\left[\mathrm{Var}(U_{ij}) + \mathrm{Var}(U_{ik})\right].$$

Consequently,

$$B_{\mathcal{T}} - A_{\mathcal{T}} = O\left[\mathrm{Var}(U_{ij}) + \mathrm{Var}(U_{ik})\right].$$

Let $\widehat{\Theta}$ denote the quasi-likelihood estimator. The asymptotic covariance matrix for $\sqrt{n}(\widehat{\Theta}-\widetilde{\Theta}_0)$ is

$$A_{\mathcal{T}}^{-1} B_{\mathcal{T}} A_{\mathcal{T}}^{-1} = A_{\mathcal{T}}^{-1} + A_{\mathcal{T}}^{-1}(B_{\mathcal{T}} - A_{\mathcal{T}})A_{\mathcal{T}}^{-1} = A_{\mathcal{T}}^{-1} + O\left[\mathrm{Var}(U_{ij}) + \mathrm{Var}(U_{ik})\right].$$

The same conclusion therefore applies to the component of the covariance matrix corresponding to $\widetilde{\beta}$. Thus, when the random-effect variances are small, the inverse information from the quasi-likelihood provides a good approximation to the sandwich variance. The Firth-like correction affects the estimator only at higher order and therefore does not alter its first-order asymptotic variance. Finally, because estimation of the taxon-specific mean relative abundances used for imputation contributes only higher-order variability to $\widehat{\beta}$ under the null $\beta = 0$, we ignore this source of variability in the variance estimation and treat the imputed mean relative abundances as fixed.

#### Real data preprocessing

The IBDMDB and MOMS-PI taxon count tables were obtained from the 16S rRNA datasets `IBD16S_mtx` and `momspi16S_mtx`, respectively, available through the `HMP2Data` R package. The ORIGINS taxon count table was obtained from the genus-level shotgun metagenomic dataset `120040_genus.biom` for Qiita study 11808 (`https://qiita.ucsd.edu/`). Because the IBDMDB and MOMS-PI datasets included repeated samples collected across visits, we restricted both datasets to samples from the first visit. ORIGINS contained one sample per participant and therefore required no visit-based filtering. We noticed that the ORIGINS taxon count table contained an unusually high proportion of singleton counts, which accounted for approximately 34% of all nonzero entries. Because singleton counts are particularly susceptible to sequencing noise and false-positive taxonomic assignments, we subtracted one from each positive count, thereby converting singleton counts to zero. For each dataset, library size was calculated before filtering as the total count across all taxa within each sample. Samples with library sizes below 5,000 were excluded.

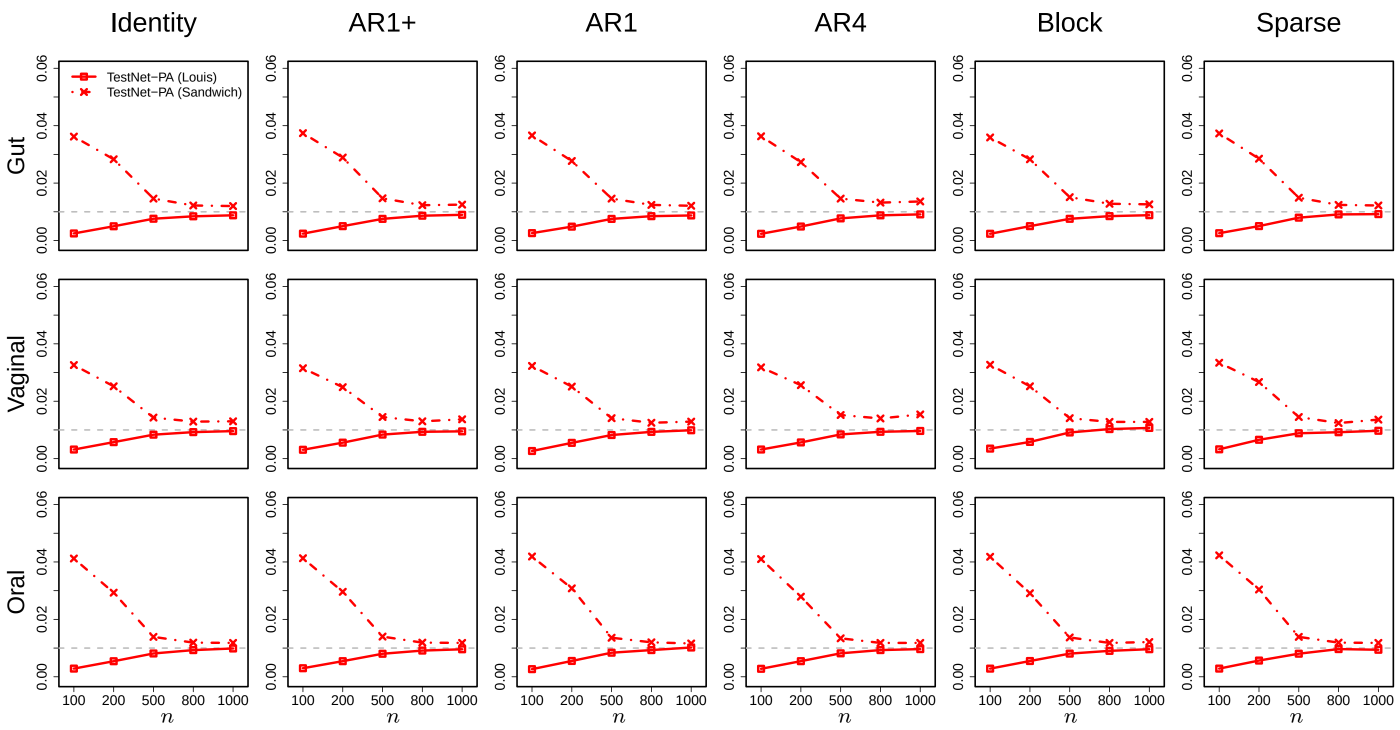


Figure S1: Simulation results for the FPR of TestNet-PA using the Louis and sandwich variance estimators, under the same settings as in Figures 2–4. The gray dashed line indicates the nominal FPR level of 1%.

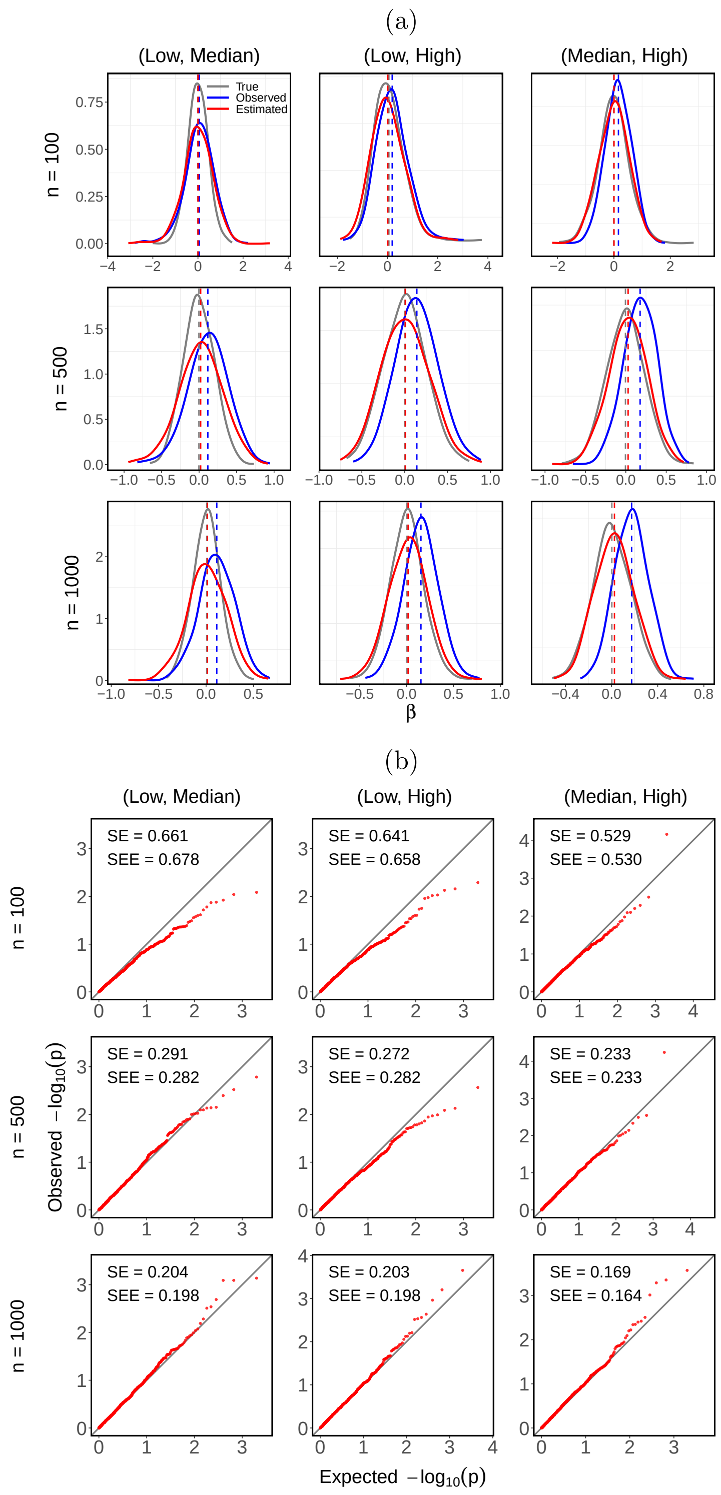


Figure S2: (a) Distributions and (b) QQ plots for the three taxon pairs comprising taxa with different prevalence levels. See more information in the caption of Figure 7.

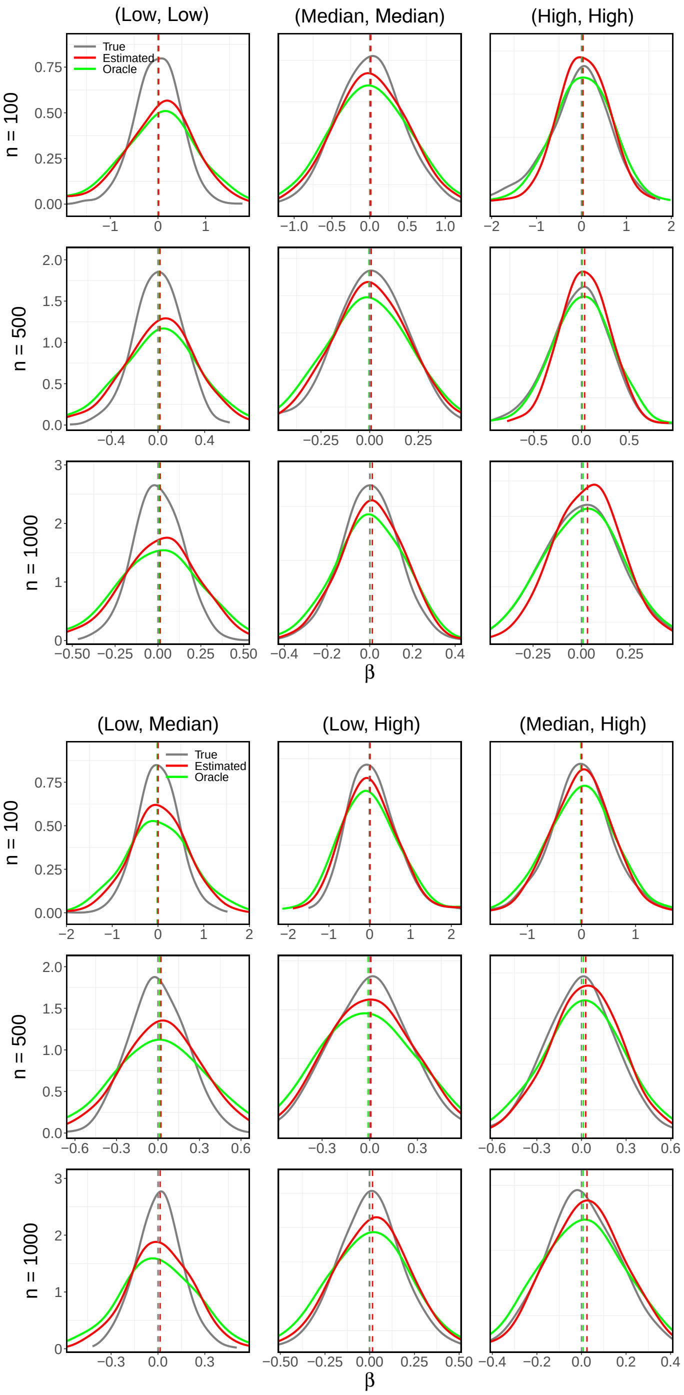


Figure S3: Distributions of the true-status-based, TestNet-PA, and oracle estimates of $\beta$ for the six taxon pairs. The true-status-based and TestNet-PA estimates are the same as those shown in Figures 7 and S2.

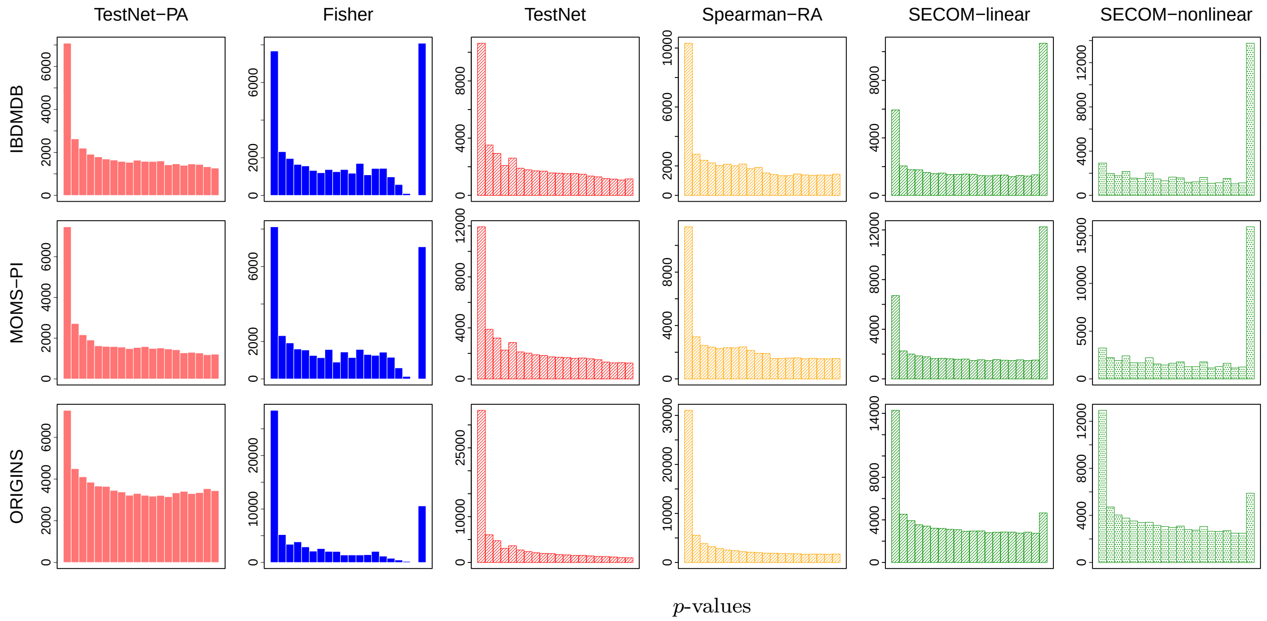


Figure S4: Distributions of $p$-values from different methods applied to the three real datasets. SparCC does not produce $p$-values and is therefore omitted.

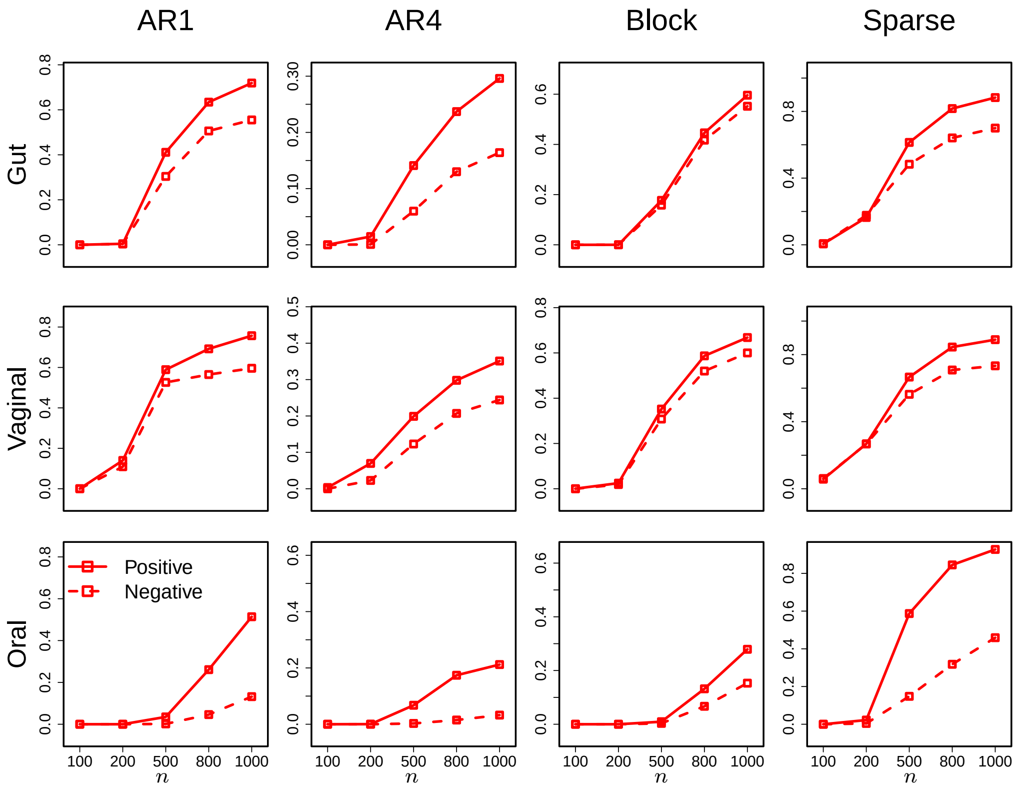


Figure S5: Simulation results comparing the sensitivity of TestNet-PA for positive versus negative dependencies. The AR1+ setting is omitted because all of its dependencies are positive.
